# Integrative analysis of a large real-world cohort of small cell lung cancer identifies distinct genetic subtypes and insights into histological transformation

**DOI:** 10.1101/2022.07.27.501738

**Authors:** Smruthy Sivakumar, Jay A. Moore, Meagan Montesion, Radwa Sharaf, Douglas I. Lin, Zoe Fleishmann, Ericka M. Ebot, Justin Newberg, Jennifer M. Mills, Priti S. Hegde, Garrett M. Frampton, Julien Sage, Christine M. Lovly

## Abstract

Small cell lung cancer (SCLC) is a recalcitrant neuroendocrine carcinoma with dismal survival outcomes. A major barrier in the field has been the relative paucity of human tumors studied. Here we provide an integrated analysis of 3,600 “real-world” SCLC cases. This large cohort allowed us to identify new recurrent alterations and new genetic subtypes, including *STK11-*mutant tumors (1.7%) and *TP53/RB1* wild-type tumors (5.5%), of which 12.7% were human papillomavirus-positive. In our cohort, gene amplifications on 4q12 are associated with increased overall survival while *CCNE1* amplification is associated with decreased overall survival. We also identify more frequent alterations in the *PTEN* pathway in brain metastases. Finally, profiling cases of SCLC containing oncogenic drivers typically associated with NSCLC demonstrates that SCLC transformation may occur across multiple distinct molecular cohorts of NSCLC. These novel and unsuspected genetic features of SCLC may help personalize treatment approaches for this fatal form of cancer.

**STATEMENT OF SIGNIFICANCE:** Minimal changes in therapy and survival outcomes have occurred in SCLC for the past four decades. The identification of new genetic subtypes, novel recurrent mutations, and an improved understanding of the mechanisms of transformation to SCLC from NSCLC may guide the development of personalized therapies for subsets of patients with SCLC.

## INTRODUCTION

Small cell lung cancer (SCLC) is the most fatal type of lung cancer, with a 5-year overall survival of ∼6%. It is estimated that SCLC kills 200,000 to 250,000 patients every year worldwide. Tobacco exposure has been linked to SCLC pathogenesis. SCLC tumors are characterized by fast growth and rapid metastatic spread to multiple sites, as well as a striking resistance to a variety of therapies (1). Patients with SCLC have not yet benefited from advances in targeted therapies and improvements from addition of immune checkpoint inhibitor (ICI) therapy have been modest (2, 3). This is in stark contrast with non-small cell lung cancer (NSCLC), where ICI therapy and targeted therapies have revolutionized treatment of molecularly defined tumor subtypes, resulting in striking increases in patient survival (4).

A major barrier towards advancing treatment paradigms for patients with SCLC has been the limited availability of tumor samples for detailed molecular characterization. In part, this is because surgical resection is uncommon, thereby limiting access to samples for analysis (5, 6). Available tumor biopsies are often small and necrotic, and rebiopsy at the time of disease progression is not standard of care. The largest published study thus far included genome sequencing of 110 SCLC genomes from resected tumors (7). Combining this study and other smaller studies with more limited genomic analyses (e.g., whole-exome sequencing and targeted sequencing such as in (8-14)), it may be possible to get an estimate of genetic alterations in a few hundred patients at best. In addition, the disparate platforms used in these different studies makes it difficult to analyze the data as a group. Moreover, there is a bias towards early-stage tumors in many of these studies. This limits our understanding of the genetic underpinning of tumor progression and metastatic spread, even though the vast majority of patients die with metastatic disease (15). One exciting new opportunity arose in the past few years with the realization that patients with SCLC often have a high number of circulating tumor cells (CTCs) (16). It is likely that longitudinal genomic analyses of SCLC CTCs will be informative regarding the mechanisms of SCLC progression, metastasis, and resistance to treatment (17). However, these studies have so far been limited to small number of patients at large medical centers capable of serially collecting and purifying CTCs. Furthermore, there is still the possibility that CTCs do not exactly reflect the biology and the genetics of primary tumors and metastases. Overall, our understanding of recurrent genetic alterations, their co-occurrence and mutual exclusivity, and how the genetic landscape of SCLC changes with tumor progression and resistance to treatment, remains limited.

Performing repeat biopsies to study molecular mechanisms of acquired resistance to tyrosine kinase inhibitors (TKIs) in *EGFR*-mutant non-small cell lung cancer (NSCLC) has been a cornerstone for the development of next-generation treatment strategies. Analysis of repeat biopsies has elucidated that histologic transformation to SCLC is detected in ∼3-14% of patients with acquired resistance to EGFR TKI therapy (18-22). These cases likely represent subclonal evolution from the original *EGFR*-mutant clonal population and not second primary cancers as they typically maintain the original *EGFR* mutation (22, 23). Histological transformation to SCLC has been repeatedly observed across multiple cohorts after emergence of resistance to first, second, and third generation EGFR TKIs. These lung adenocarcinomas show loss of *RB1* as they transition to the SCLC phenotype (21, 24) and evidence from a few cases suggests that mutation signatures and CNVs may also change during this transition (25). Other data suggest that SCLC transformation from lung adenocarcinoma is mainly driven by transcriptional reprogramming rather than genetic events (26). Still, despite the clinical importance of SCLC transformation from NSCLC, our understanding of this transformation process and its genetic basis remains incomplete.

To address some of the limitations of previous studies, we evaluated a large cohort of real-world SCLC cases. This cohort of 3,600 cases is more than 30 times larger than the largest published study (7) and allowed us to identify new genetic subgroups in SCLC, site-specific mutational patterns, and insights into histological transformation from NSCLC. The new insights gained from this integrated analysis open new avenues of research in the field and readily suggest new therapeutic opportunities for subsets of patients with SCLC.

## RESULTS

### Clinical characteristics of a large cohort of real-world patients with SCLC

The cohort consisted of 3,600 patients with SCLC biopsy samples submitted to Foundation Medicine, Inc. for tumor genomic profiling (**Fig. 1A** and **Table 1A**). 52.1% of patients were female and 47.9% were male, with a median age of 65 years (range: 21-89 years). 3,114 patients (85.5%) were predicted to have a European ancestry (EUR), with the remaining patients classified into the following ancestry groups: African (AFR, n= 256, 7.1%), Admixed American (n=150, 4.2%), East Asian (EAS, n=69, 1.9%), South Asian (n=11, 0.3%). Tumor samples were most frequently profiled from lung (37.8%), but also comprised other sites such as regional/distant lymph nodes (n=901, 25.0%), liver (n=747, 20.8%), brain (n=116, 3.2%) and soft tissue (n=114, 3.2%). As expected from previous analyses (7), most cases (99.5%) were microsatellite stable. Tumor mutational burden (TMB) varied across cases (range: 0 – 276.3 mutations/Megabase; mut/Mb), with a median of 7.8 mut/Mb and a mean of 9.5 mut/Mb; 38.9% cases had a TMB-high status (TMB≥10 mut/Mb). TMB was similar across patients from diverse genetic ancestry (**Supplementary Fig. S1**).

**Figure 1:**
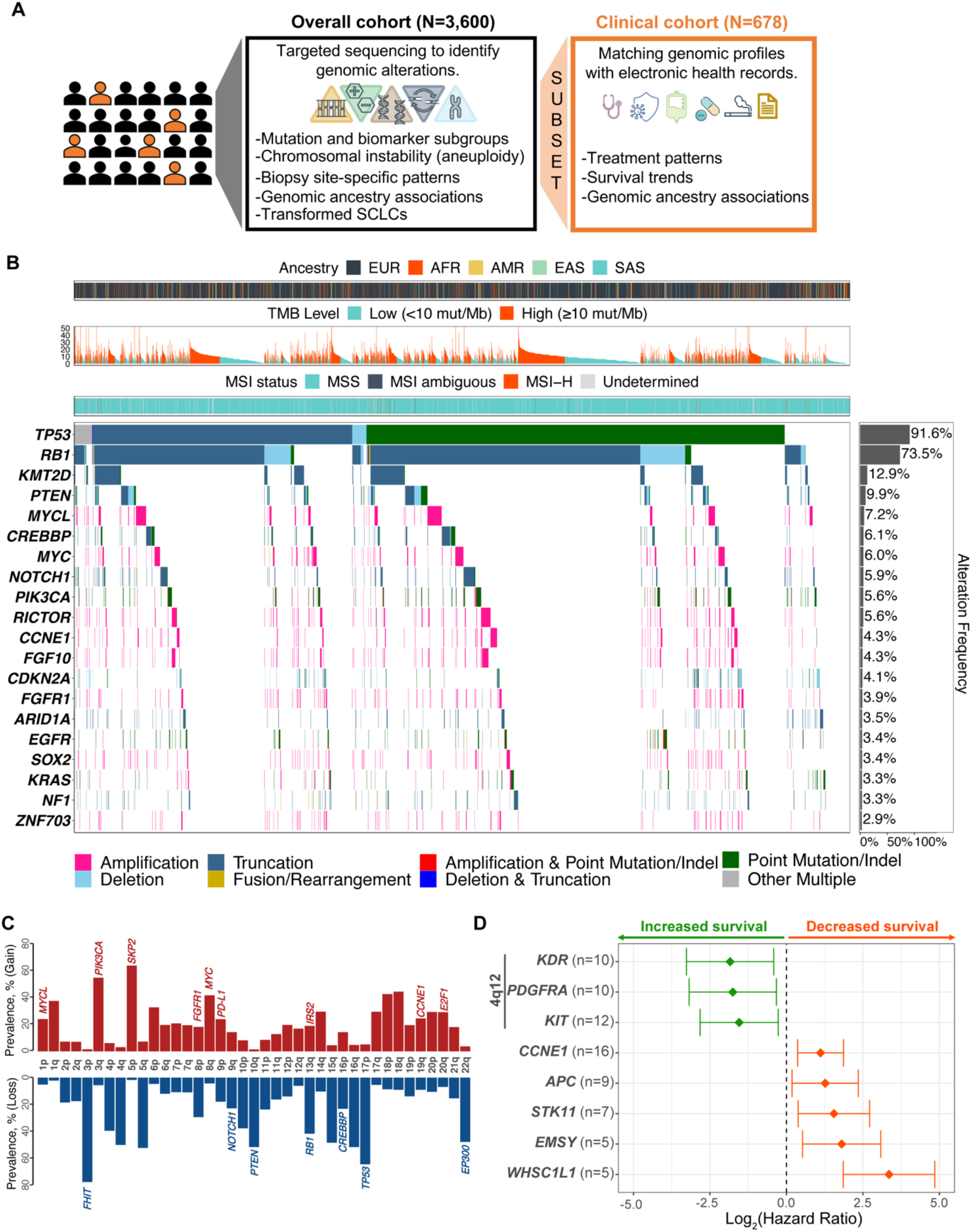
The genomic landscape of SCLC tumors in a real-world cohort of patients with SCLC. **A**. Schematic representation of the overall study design in the SCLC cohort comprising 3,600 patients, including 678 cases for which clinical data are available. **B**. Patterns of the 20 most frequent gene alterations identified in SCLC tumors. Genes are indicated on the left and their alteration frequency on the right. Predicted genomic ancestry, tumor mutational burden (TMB) and microsatellite instability (MSI) status for each patient is overlayed on top. **C**. Prevalence of chromosomal copy number loss (blue) and gain (red) in SCLC tumors. Notable genes are indicated for some chromosome arms. **D**. Association between overall survival and genetic alterations (for genes altered in ≥5 cases) in the clinical cohort of evaluable stage III/IV SCLC (N=511). Genes identified to be associated with increased survival are shown in green and those identified to be associated with decreased survival are shown in orange (P≤0.05). *KDR*, *PDGFRA*, and *KIT* are all located on 4q12. After FDR-based adjustment, only *WHSC1L1* was statistically significant (adj P=0.001).

**Table 1:**
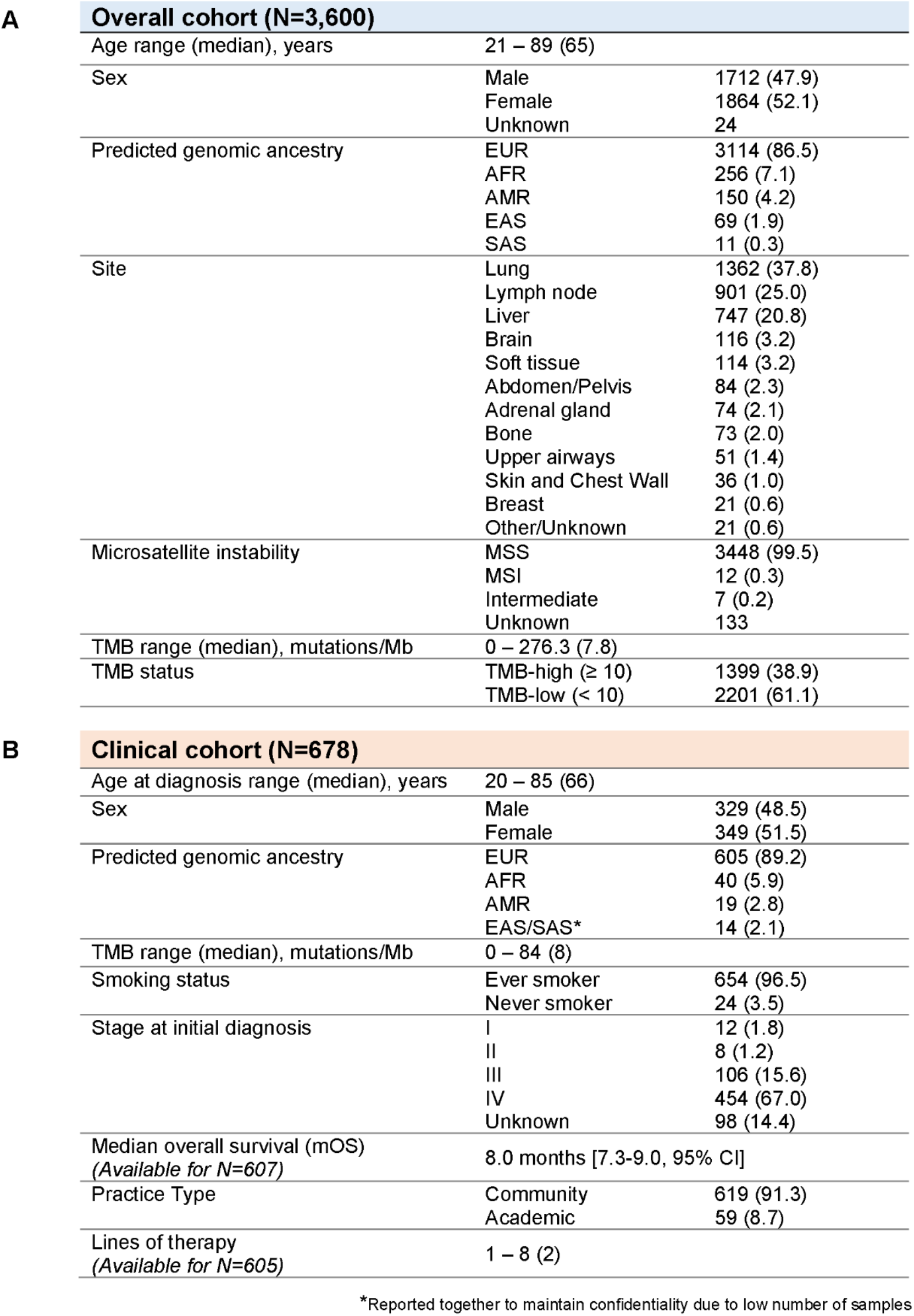
Clinicopathological characteristics of the real-world small cell lung cancer cohort. **A.** Clinicopathological characteristics of 3,600 patients (overall cohort) as well as **B**. 678 patients with clinical information (clinical cohort). The clinical cohort had a large, but not complete, overlap with the overall cohort.

|  |  |  |  |
| --- | --- | --- | --- |
| **A** | **Overall cohort (N=3,600)** |  |  |
|  | Age range (median), years | 21 – 89 (65) |  |
|  | Sex | Male | 1712 (47.9) |
|  |  | Female | 1864 (52.1) |
|  |  | Unknown | 24 |
|  | Predicted genomic ancestry | EUR | 3114 (86.5) |
|  |  | AFR | 256 (7.1) |
|  |  | AMR | 150 (4.2) |
|  |  | EAS | 69 (1.9) |
|  |  | SAS | 11 (0.3) |
|  | Site | Lung | 1362 (37.8) |
|  |  | Lymph node | 901 (25.0) |
|  |  | Liver | 747 (20.8) |
|  |  | Brain | 116 (3.2) |
|  |  | Soft tissue | 114 (3.2) |
|  |  | Abdomen/Pelvis | 84 (2.3) |
|  |  | Adrenal gland | 74 (2.1) |
|  |  | Bone | 73 (2.0) |
|  |  | Upper airways | 51 (1.4) |
|  |  | Skin and Chest Wall | 36 (1.0) |
|  |  | Breast | 21 (0.6) |
|  |  | Other/Unknown | 21 (0.6) |
|  | Microsatellite instability | MSS | 3448 (99.5) |
|  |  | MSI | 12 (0.3) |
|  |  | Intermediate | 7 (0.2) |
|  |  | Unknown | 133 |
|  | TMB range (median), mutations/Mb | 0 – 276.3 (7.8) |  |
| | TMB status | TMB-high ($\geq 10$) | 1399 (38.9) |
| | | TMB-low ($< 10$) | 2201 (61.1) |
| **B** | **Clinical cohort (N=678)** |  |  |
|  | Age at diagnosis range (median), years | 20 – 85 (66) |  |
|  | Sex | Male | 329 (48.5) |
|  |  | Female | 349 (51.5) |
|  | Predicted genomic ancestry | EUR | 605 (89.2) |
|  |  | AFR | 40 (5.9) |
|  |  | AMR | 19 (2.8) |
|  |  | EAS/SAS* | 14 (2.1) |
|  | TMB range (median), mutations/Mb | 0 – 84 (8) |  |
|  | Smoking status | Ever smoker | 654 (96.5) |
|  |  | Never smoker | 24 (3.5) |
|  | Stage at initial diagnosis | I | 12 (1.8) |
|  |  | II | 8 (1.2) |
|  |  | III | 106 (15.6) |
|  |  | IV | 454 (67.0) |
|  |  | Unknown | 98 (14.4) |
|  | Median overall survival (mOS)<br>(Available for N=607) | 8.0 months [7.3-9.0, 95% CI] |  |
|  | Practice Type | Community | 619 (91.3) |
|  |  | Academic | 59 (8.7) |
|  | Lines of therapy<br>(Available for N=605) | 1 – 8 (2) |  |

\*Reported together to maintain confidentiality due to low number of samples

Of the 3600 patients, 678 patients had additional clinical data derived from electronic health records (EHR) as part of the clinico-genomic database (CGDB) (**Table 1B**). Characteristics of the CGDB cohort were analogous to the overall dataset, with median age of 66 years (range: 20-85 years), a similar distribution based on sex (51.5% female, 48.5% male), predominantly EUR ancestry (89.2%) and similar TMB distributions (median: 8 mut/Mb, range: 0-84 mut/Mb). This sub-cohort also contained additional clinico-pathological and treatment data. Most patients had a smoking history (96.5%) and exhibited advanced stage disease at the time of initial diagnosis. As expected, 91.3% of patients in CGDB were treated in community setting compared to 8.7% treated in an academic setting. Median overall survival (OS) for the cohort was 8.0 months (95% CI, 7.3 – 9.0 months). Patients received a median of two lines of therapy (LoT, range: 1-8) prior to tumor genomic profiling. 86.2% of patients received first line platinum-based doublet chemotherapy, and 23.3% of patients received a PD-L1 inhibitor in the first line (**Supplementary Table S1**). In the second line and beyond, patients were treated with multiple different chemotherapies and immune checkpoint inhibitors, with no one therapy representing a majority, reflecting the clinical uncertainty of how to treat relapsed SCLC.

Utilizing this large dataset, we sought to perform an integrative analysis of SCLC tumors to gain a better understanding of the genetic underpinning of SCLC progression and metastasis.

### Overview of recurrent genomic alterations in 3,600 cases of SCLC

Deep sequencing of exons from 324 cancer-related genes and select introns from up to 34 genes frequently rearranged in cancer (**Supplementary Table S2**) revealed a number of recurrent alterations. As expected, alterations resulting in the inactivation of the *TP53* and *RB1* genes were the most frequent and observed in 91.6% and 73.5% cases, respectively (**Fig. 1B** and **Supplementary Table S3**). In *TP53*–mutant samples inactivation was observed mostly through short variants, comprising base substitutions and indels (98%), whereas *RB1-*mutant cases included high rates of short variants (85%) and focal copy number loss (14%) (**Supplementary Table S3**). The base substitutions were similar to what has been observed in NSCLC, where carcinogens from cigarette smoke are also frequent drivers of cancer initiation (**Supplementary Fig. S2**) (27).

Our analysis also confirmed the previously described recurrent loss-of-function alterations in *KMT2D* (*MLL2*) (12.9%), *CREBBP* (6.1%), and *NOTCH1* (5.9%) and gain-of-function events and copy number amplifications in *MYC* (6.0%), *MYCL* (7.2%), and *SOX2* (3.4%) (7-9) (**Fig. 1B,C** and **Supplementary Table S3**). Compared to previous smaller-scale studies, we noted an increased representation of alterations in PI3K pathway genes (e.g., *PTEN* 9.9%, *PIK3CA* 5.6%, *RICTOR* 5.6%) and the RAS/MAPK pathway genes (e.g., *EGFR* 3.4%, *KRAS* 3.3.%, *NF1* 3.3%) (see below) (**Fig. 1B,C** and **Supplementary Table S3**). We observed similar trends through analysis of cell-free DNA (cfDNA) obtained from peripheral blood liquid biopsies from 81 patients with SCLC (**Supplementary Fig. S3A** and **Supplementary Table S4**), which is abundant in SCLC compared to NSCLC and other tumor types (**Supplementary Fig. S3B**) (28).

At the chromosomal level, loss of chromosome arms 3p and 17p were the most common (77.6% and 64.5% respectively) in SCLC tumors, whereas chromosome arm-level gains were frequently observed in 5p and 3q (64.0% and 55.0% respectively), among other regions (**Fig. 1C** and **Supplementary Table S5**). Overall, chromosome arm losses were more frequent than gains in SCLC tumors, and these losses were significantly enriched in regions with a high tumor suppressor gene score (29) (**Fig. 1C**, **Supplementary Fig. S4**, and **Supplementary Table S5**), providing additional evidence that SCLC is in large part driven by loss of tumor suppressors.

### Novel recurrent gene mutations in SCLC

Through our analysis of such a large dataset, we were able to identify genes for which mutations have not been previously associated with SCLC (**Supplementary Table S3**). For example, while alterations in *KEAP1* have mostly been associated with NSCLC (30-32), ∼3% of SCLC samples had mutations in this gene, suggesting that inactivation of *KEAP1* may contribute to SCLC pathogenesis. Mutations in *TET2*, which contribute to the development of hematologic malignancies (33), are also found in ∼2% of SCLC cases, indicating again of a possible tumor suppressor role in SCLC. As a final example, recurrent amplification events for *ZNF703* (2.9%) supports that the transcriptional regulator coded by the gene has oncogenic potential in SCLC, possibly because ZNF703 can control the expression of SOX2 (34), a known oncogene in SCLC (9) (**Fig. 1B** and Supplementary **Table S3**).

### Recurrent gene rearrangements in SCLC

We identified frequent gene rearrangements in 338 tumors, including in *RB1* (n=31), *NOTCH1* (n=11), *CREBBP* (n=10), *KMT2D* (n=9), and *TP53* (n=8) suggesting that these events contributed to inactivation of these tumor suppressors (**Supplementary Fig. S5A**). This analysis also identified several rearrangements not previously described in SCLC, including events involving *ETV6* (n=15). Based on the oncogenic role of rearrangements involving the transcription factor coded by this gene in leukemia and solid tumors (35, 36), these observations suggest that *ETV6* rearrangements could also contribute to SCLC development. Intriguingly, tumors with gene rearrangements showed enrichment for specific other genetic alterations, including mutations in *MCL1*, which codes for an anti-apoptotic factor, as well as in *SMARCA4* and *DNMT3A*, which code for epigenetic modulators (**Supplementary Fig. S5B and Supplementary Table S6).**

### Gene amplifications on 4q12 are associated with increased overall survival in patients with SCLC

The availability of a large group of patients with both genetic characterization and clinical data provided us with a unique ability to determine if any of the recurrently detected genomic alterations were associated with median overall survival (mOS). For the entire clinical cohort with survival information (n=607), the mOS from the date of initial diagnosis was 8.0 months [7.2 – 9.0, 95% CI] across patients of different genetic ancestry (**Supplementary Fig. S6A**) and varied by stage of initial diagnosis (**Supplementary Fig. S6B**).

We focused our analysis on patients with advanced disease (stage III/IV SCLC, N=511). We identified recurrent gene amplification events at 4q12 associated with increased survival (**Fig. 1E** and **Supplementary Table S7**). This region contains three genes coding for receptor tyrosine kinases (RTKs), *KDR* (coding for VEGFR2), *PDGFRA*, and *KIT*. Alterations in these genes were associated with improved OS with a mOS of 67.9 months (*KDR*, *PDGFRA*) and 24.0 months (*KIT*) (**Supplementary Table S7**). c-KIT has been investigated unsuccessfully as a possible drug target in SCLC (37), based in part on evidence that high levels of c-KIT can be associated with worse overall survival (38, 39). However, other studies suggest that low protein expression of c-KIT can be correlated with worse survival (40), and no study has examined co-expression of the three RTKs, and it is possible that too much RTK signaling upon genomic amplification might slow the expansion of SCLC tumors. Of note, amplifications on 4q12 were observed in 1.1% of the overall SCLC cohort. We also identified genes whose recurrent inactivation is associated with worse survival, including mutations in *APC*, which codes for a Wnt pathway regulator and whose loss has been associated with relapsed SCLC (12), as well as a new association between amplification of the *CCNE1* gene, which codes for the cell cycle regulator Cyclin E1, and worse survival in patients with SCLC (**Fig. 1E** and **Supplementary Table S7**). Together, these analyses identify new candidate factors that may play a role in the development and the progression of SCLC.

### Mutation Spectrum by Different Sites of Metastasis

Clinical prognostic factors for patients with SCLC include stage and extent of disease (**Table 1**). Overall, the number of organ sites involved is inversely related to prognosis, with certain metastatic sites – such as CNS, liver, and bone marrow – conferring poorer prognosis (14,41,42). However, the genetic underpinning of SCLC metastasis is largely unknown (15). We examined if tumors at different sites had distinct patterns of genetic alterations, focusing on major sites of metastasis, and grouping more rare sites by general area in the body (**Table 1**).

When we first examined tumor site and TMB, we made the unexpected observation that brain metastases had the highest TMB (median TMB 10.0 mut/Mb), with adrenal gland metastases (median TMB 9.6 mut/Mb) also having a significantly higher TMB compared to lung tumors (median TMB 7.8 mut/Mb) (**Fig. 2A**). Of note, brain metastases also had a higher prevalence of TMB-high samples, defined at a threshold of 10 mut/Mb (50.9% vs. 38.7%, P = 0.01, **Supplementary Table S8**). Brain and liver metastases also showed a significant increase in chromosomal arm-levels gains compared to lung-biopsied tumors (**Supplementary Fig. S7A** and **Supplementary Table S9**), further suggesting unique genetic mechanisms. We and others have identified *Nfib* amplification in metastatic mouse SCLC tumors (43-45). While NFIB expression is high in a large fraction of SCLC metastasis (44, 46), the *NFIB* gene is rarely amplified in human SCLC tumors (7), even though this amplification may be selected in human SCLC cell lines (47). The *NFIB* gene itself is not analyzed in our panel, but we did not find significant gain of 9p where *NFIB* is located (**Supplementary Table S9**). The identity of potential drivers of SCLC metastasis on chromosome 16p, the top gain (**Supplementary Fig. S7B**), remains unknown, but genomic gain of 16p13.3 has been associated with poor outcome in prostate cancer (48) and this region contains the *PDK1* gene, coding for a component of the PI3K/AKT pathway.

**Figure 2:**
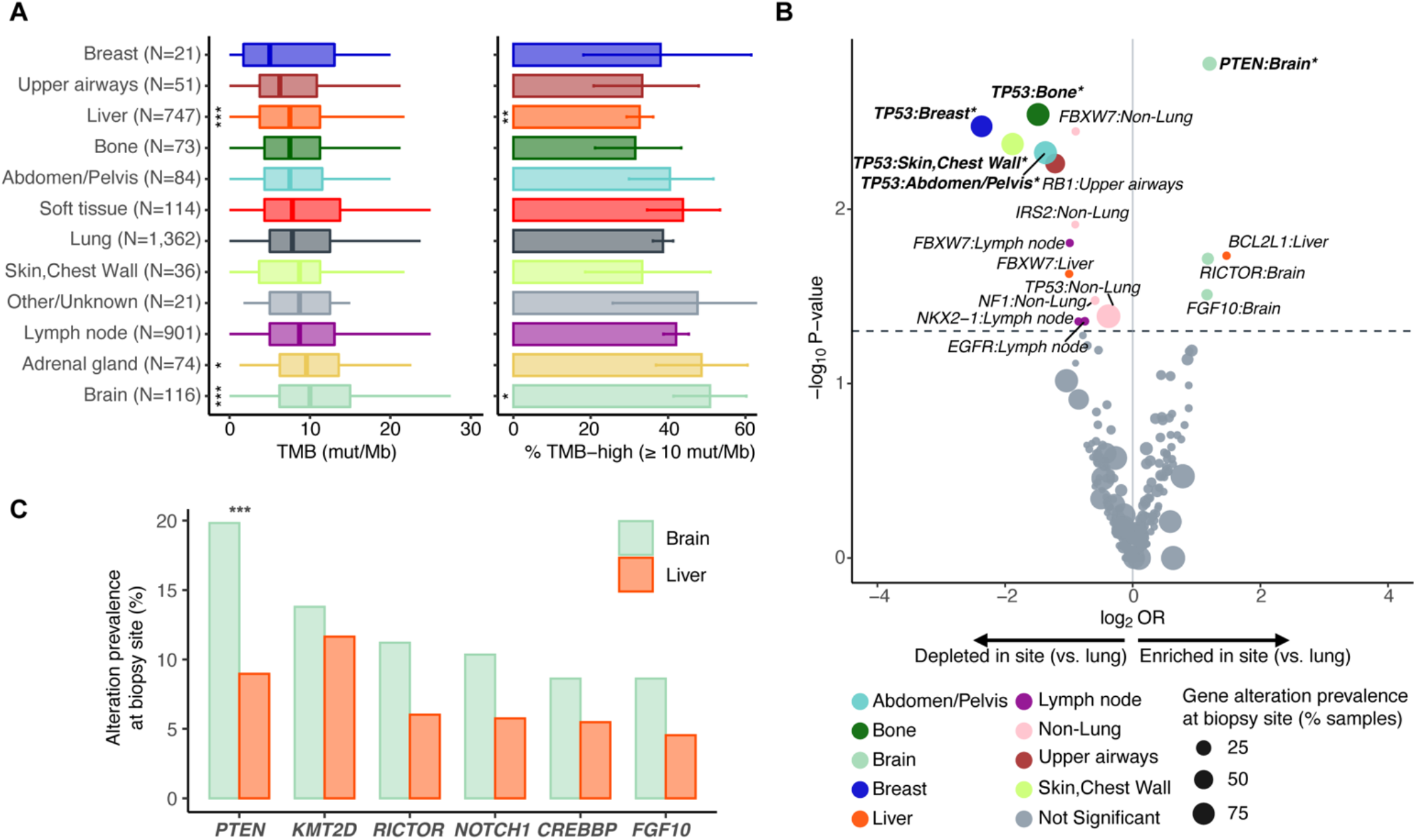
Biopsy-site specific patterns of tumor mutational burden and gene alterations in SCLC. **A**. Patterns of tumor mutation burden (TMB) in SCLC tumors by anatomical location. Shown are boxplots of the TMB distribution (left) and the percentage of TMB-high samples (right) at each site. The sites are ordered by their median TMB from lowest (top) to highest (bottom). **B**. Volcano plot showing patterns of enrichment or depletion of recurrent gene alterations in different tumor sites compared to lung biopsied-SCLC tumors. Gene alterations identified to be statistically different in prevalence between a metastatic site and lung are indicated, with those statistically significant after correcting for multiple hypothesis testing in bold. **C**. Prevalence and comparison of the most frequently identified gene alterations in brain *vs.* liver SCLC metastases. (P value thresholds: *: 0.05, ***: 0.001)

Importantly, when we investigated alterations in specific genes, we found that brain metastases were significantly enriched for *PTEN* alterations compared to lung tumors (19.8% vs. 9.7%, P=0.012) (**Fig. 2B** and **Supplementary Table S10**). This was also true for brain vs. liver metastases (19.8% vs. 9.0%, P=0.007) (**Fig. 2C** and **Supplementary Table S11**). Similar observations were made for *RICTOR* amplification, a gene that codes for a regulator of mTOR activity (49), although this did not reach statistical significance after correcting for multiple testing (11.2% brain metastases vs. 5.3% lung biopsies). These data suggest that the PI3K/AKT/MTOR pathway may play a unique role in SCLC brain metastasis, thereby suggesting a new potential therapeutic vulnerability.

### Defining genetic subtypes in SCLC

A previous study of 110 SCLC tumors failed to identify clear genetic subtypes (7). In contrast, a growing consensus in the field is that SCLC subtypes may be defined by transcriptional signatures driven by specific transcription factors (50-52). Importantly, some of the subtypes may have unique vulnerabilities, including for example sensitivity to Aurora kinase inhibitors in the *MYC*-high NEUROD1-high SCLC subtype (SCLC-N) (53). The large number of tumors analyzed enabled us to investigate genetic interactions in SCLC defined by co-occurrence and mutual exclusivity that may help conclusively determine whether such genetic subtypes may exist in SCLC (**Supplementary Fig. S8** and **Supplementary Table S12**). Not surprisingly, mutations in *TP53* and *RB1* were highly significantly co-occurring (Odds ratio, OR = 7.4, p < 10^-5^). Some observations were more unexpected: for example, alterations in the *PIK3CA* gene that activate the catalytic subunit of the PI3K kinase are significantly co-occurring with amplification of the gene coding for SOX2 suggesting cooperative effects between these two oncogenes in SCLC (54). Our analysis of recurrent mutations and alterations further suggested three possible subtypes. First, some tumors had no alterations in *TP53* and/or *RB1*. Second, we noticed a subgroup of tumors with *STK11* mutations (*STK11* codes for the LKB1 kinase). Third, our analysis found SCLC tumors with oncogenic driver mutations characteristic of NSCLC, suggestive of transformation from NSCLC to SCLC. These genetic groups are further discussed below.

### SCLC tumors without genomic inactivation of *TP53* and/or *RB1*

While SCLC is viewed as a cancer type in which cells are functionally mutant for RB and p53, not all SCLC cells show inactivation of the *RB1* and *TP53* genes. For example, chromothripsis, a catastrophic event characterized by massive genomic rearrangements, has been suggested to lead to amplification of the *CCND1* gene coding for Cyclin D1, which may result in the functional inactivation of RB by increased phosphorylation (7). Importantly, SCLC cells expressing wild-type RB may be sensitive to CDK4/6 inhibitors (55). Similarly, strategies are being developed to treat p53 wild-type tumors (56). Thus, a better understanding of SCLC tumors that have functional RB and/or p53 molecules may help develop distinct targeted therapies for these patients.

We analyzed SCLC tumors in our cohort that may be *RB1* and/or *TP53* wild-type. For this analysis, we defined *RB1* and *TP53* mutant tumors as tumors where we could detect pathogenic alterations in these genes; wild-type tumors had no pathogenic variants and no variants of unknown significance (VUS). We also assumed that tumors with one detected mutant allele of *RB1* or *TP53* may have lost the other allele *via* gene silencing or other less readily detectable events (and in some cases, we are able to document this loss of heterozygosity). This analysis of 3590 tumors (excluding 10 tumors with VUS in *RB1* or *TP53*) identified 96 *TP53* wild-type tumors (2.7%), 747 *RB1* wild-type tumors (20.8%), and 197 tumors wild-type for both *TP53* and *RB1* (5.5%). Presence of wild-type *RB1* was significantly associated with a lower TMB, and tumors with both *TP53* and *RB1* wild-type genes had the lowest TMB (**Fig. 3A,B**, **Supplementary Fig. S9A,B**, and **Supplementary Fig. S10A,B**).

**Figure 3:**
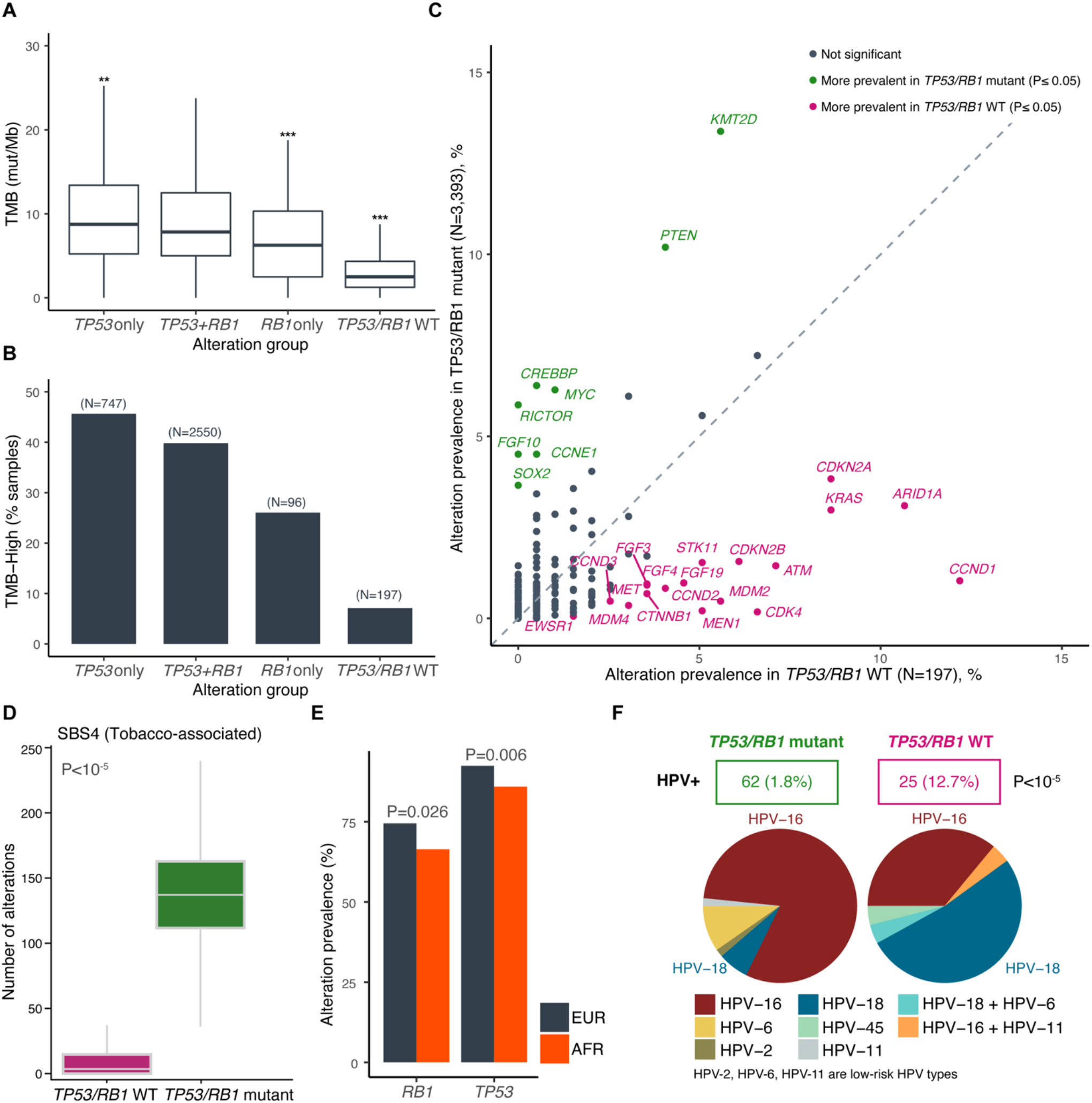
*TP53* and *RB1* wild-type tumors represent a distinct genetic subtype of SCLC associated with HPV infection. **A**. Distribution of tumor mutation burden (TMB, mutations/Mb) in SCLC tumors wild-type and/or mutant for *TP53* and *RB1*. The TMB in each mutation group was compared against cases identified to be *TP53*+*RB1* double mutants (P value thresholds: **: 0.01, ***: 0.001). **B**. Prevalence of TMB-high status (≥10 mutations/Mb) in SCLC tumors wild-type and/or mutant for *TP53* and *RB1*. **C**. Prevalence of gene alterations in SCLC tumors wild-type for *TP53* and *RB1* compared to *TP53* or *RB1* mutant SCLC tumors. **D**. Analysis of the number of mutations representing the tobacco-associated signature (SBS4) in *TP53*/*RB1* wild-type and mutant SCLC tumors. **E**. Comparative prevalence of *TP53* and *RB1* alterations in patients of European (EUR) and African (AFR) ancestry. **F**. Detection of human papillomavirus (HPV) and breakdown of HPV types in *TP53*/*RB1* mutant and wild-type SCLC tumors. HPV-16, HPV-18 and HPV-45 are high risk HPVs whereas HPV-2, HPV-6 and HPV-11 are low-risk HPVs.

We also used the large size of our cohort to examine co-occurrences and mutual exclusivity events in *TP53* and/or *RB1* wild-type tumors compared to mutant tumors (**Supplementary Table S13**). As expected, inactivating alterations in *CDKN2A*, which codes for p16, a positive regulator of RB, and activation alterations in *CCND1*, which codes for Cyclin D1, a negative regulator of RB, were significantly mutually exclusive with alterations in *RB1*. Similarly, amplification of *MDM2*, a negative regulator of p53, was significantly mutually exclusive with inactivation of *TP53* (**Fig. 3C**, **Supplementary Fig. S9C**, and **Supplementary Fig. S10C**). While *CCNE1* amplification events are frequent in SCLC tumors, and while Cyclin E/CDK2 kinase complexes inhibit RB function, *CCNE1* amplification was not enriched in *RB1* wild-type tumors (**Supplementary Fig. S10C**). Because *CCNE1* amplification was enriched in *TP53* mutant tumors (**Supplementary Fig. S10C**), it is possible that p53 loss is important to prevent cell death or cell cycle arrest upon increased Cyclin E/CDK2 activity.

Enrichment in *TP53* and/or *RB1* wild-type tumors in alterations in genes such as *FGFR1*, *KRAS*, *KEAP1*, or *BRAF* (**Fig. 3C, Supplementary Fig. S9C**, and **Supplementary Fig. S10C**), which are frequently mutated in NSCLC, suggest that a number of these tumors may have arisen from NSCLC (see below). In support of this possibility, *TP53* and *RB1* wild-type tumors had a lower signature associated with smoking (**Fig. 3D** and **Supplementary Fig. S11**), and *TP53* and *RB1* alterations were less frequent among the 24 never-smokers available in the clinical cohort (*TP53* 96.5% ever-smokers vs. 45% never-smokers, *RB1* 77.0% vs. 45%; **Supplementary Fig. S12**).

There was no significant difference in median OS (mOS) between *TP53*/*RB1* mutant and wild-type tumors (**Supplementary Fig. S13A**), supporting the notion that the wild-type tumors were indeed SCLC. When assessed separately, *RB1* wild-type and mutant tumors also showed no differences in mOS (**Supplementary Fig. S13B**). *TP53*-mutant tumors showed a slightly lower mOS compared to *TP53* wild-type tumors (8.0 vs. 8.8 months, HR = 1.6 [1.1-2.5], P = 0.03; **Supplementary Fig. S13C**), although this analysis is limited by the small cohort size of *TP53* wild-type tumors.

Notably, AFR genetic ancestry was associated with decreased presence of *TP53* or *RB1* alterations (**Fig. 3E**). Further, when examining alterations in young patients of all genetic ancestries (<50y; N=239) compared to older patients (≥50y; N=3,361), *TP53* and *RB1* alterations were less frequent in younger patients: 77.0% vs. 92.6% *TP53* (P<10^-4^), 60.7% vs. 74.4% *RB1* (P<10^-4^), respectively (**Supplementary Fig. S14** and **Supplementary Table S14**).

### Identification of human papilloma virus (HPV) in SCLC

The presence of tumors with no genetic alterations in *TP53* and *RB1* led us to investigate if the p53 and RB proteins may be inactivated by other means in these tumors. Oncoproteins for several human viruses can functionally inactivate p53 and RB (e.g., E6/E7 from human papillomavirus, HPV) (57, 58). Sequencing reads left unmapped to the human reference genome were compared against strains of oncoviruses, as described previously (59). This analysis identified 87 tumors with HPV sequences.

Strikingly, 12.7% (25/197) *RB1/TP53* wild-type tumors were HPV-positive, while only 1.8% (62/3,393) of *RB1/TP53* mutant tumors were HPV-positive (**Fig. 3F**). Most of the HPV-positive *RB1/TP53* wild-type cases were HPV16/18/45, which are among the most oncogenic subtypes. We also noted six tumors positive for the Merkel cell polyomavirus, all of which were *RB1/TP53* mutant tumors; this small number made it difficult to draw any conclusions.

Taken together, these data indicate that tumors with wild-type *RB1* and/or *TP53* define a distinct subgroup of SCLC tumors that may be amenable to unique treatment modalities.

### Cohort of SCLC tumors defined by *STK11* mutations

Genetic alterations in *STK11* are frequent in NSCLC. Functional inactivation of LKB1 has been shown to promote tumor development and metastasis (60, 61) and modify response to ICI therapy (62-64). Recurrent *STK11* mutations are also found in a subset of NSCLC-like large-cell neuroendocrine cancers (LCNECs) (31). In contrast, *STK11* mutations have not been identified as recurrent events in SCLC, and little is known about LKB1 in SCLC (65). We identified 62 *STK11*-mutant tumors in our cohort, representing 1.7% of the entire cohort (**Fig. 4A**). These tumors were enriched for mutations in genes associated with the development of NSCLC, including *KRAS* and *KEAP1*, and harbored fewer mutations in *RB1* compared to the *STK11*-wild-type cohort (**Fig. 4A,B, Supplementary Table S15**).

**Figure 4:**
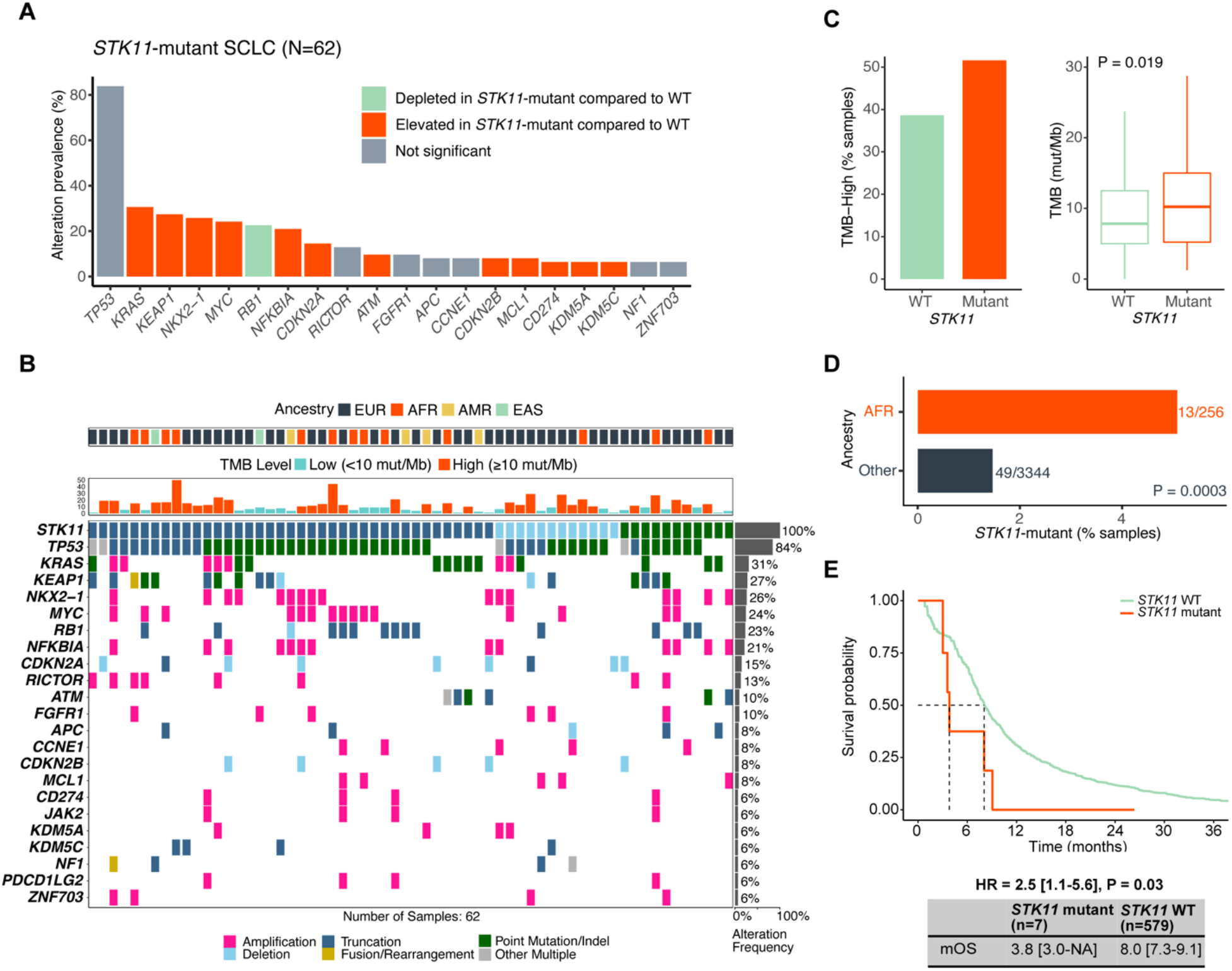
*STK11* alterations define a new genetic subtype of SCLC with decreased overall survival. **A**. Most frequently mutated gene alterations in *STK11*-mutant SCLC tumors. Gene alterations identified more frequently in *STK11*-mutant SCLC tumors compared to *STK11* wild-type tumors are displayed in mint green and those identified less frequently in *STK11*-mutant SCLC tumors compared to *STK11* wild-type tumors in orange. Genes that were similarly prevalent are shown in grey. **B**. Oncoplot of the most frequent gene alterations identified in *STK11*-mutant SCLC tumors. Genes are indicated on the left and their alteration frequency on the right. Ancestry and tumor mutation burden (TMB) for each case, are indicated on top. **C**. Distribution of tumor mutation burden (left) and the prevalence of TMB-high status, defined at ≥10 mutations/Mb (right), in *STK11*-mutant and wild-type SCLC tumors. **D**. Comparative prevalence of *STK11* alterations in patients of African (AFR) ancestry compared to other ancestry groups (NA: not available due to low sample counts). **E**. Overall survival of patients with *STK11* mutant and wild-type SCLC tumors. Samples with unknown/ambiguous functional status or reduced quality were excluded from the *STK11* wild-type cohort.

These tumors also showed a significant trend for higher TMB (**Fig. 4C**). Patients with *STK11*-mutant SCLC tended to be enriched for AFR origin (**Fig. 4D**). Although the number of patients with *STK11*-mutant tumors was limited in the clinical cohort (n=7), we observed a reduction in survival in these patients (**Fig. 4E**). These findings define a rare subtype of SCLC tumors (1-2%) with mutant *STK11*. Patients with *STK11*-mutant SCLC tumors may benefit from efforts in other cancer types with the same mutations to develop new therapeutic approaches (66).

### NSCLC driver mutations are recurrently detected in SCLC, suggesting that SCLC transformation is more frequent than suspected

Histological transformation from NSCLC to SCLC has been observed in ∼3-14% of patients with *EGFR*-mutant NSCLC that develop acquired resistance to *EGFR* tyrosine kinase inhibitor (TKI) therapy (21,22,67). These SCLC tumors are thought to represent subclonal evolution from the original *EGFR*-mutant clonal population and not a new primary tumor, as they maintain the original *EGFR* mutation (22, 24). Previous studies have suggested that *EGFR*-mutant transformed SCLC (t-SCLC) may adopt similar genomic and phenotypic characteristics of *de novo* SCLC but have been limited by very small sample sizes.

In our cohort, we identified 107 SCLC samples harboring activating mutations within the *EGFR* kinase domain. The genetic ancestry of this subgroup of patients was composed of 71.0% EUR (n=76), 8.4% AFR (n=9), 3.7% AMR (n=4), 15.0% EAS (n=16), and 1.9% SAS (n=2), suggesting that lineage transformation can occur across all genomic ancestries. Overall, the *EGFR*-mutant t-SCLC cohort mirrored the genomic landscape of *de novo* SCLC, with *RB1* and *TP53* mutations in the majority of sample (95.3% and 83.1% respectively; **Fig. 5A** and **Supplementary Table S16**). Notably, *PIK3CA* mutations were enriched in the *EGFR*-mutant t-SCLC cohort compared to SCLC samples lacking an *EGFR* alteration (25.2% vs. 5.0%, OR 6.4, P<10^-5^). Other genetic events enriched in *EGFR*-mutant t-SCLC included amplification and gain of function alterations in *NFKBIA* (10.3% vs. 1.6%), *NKX2-1* (13.1% vs. 2.5%) and *CCNE1* (10.3% vs. 4.1%), *RBM10* loss of function mutations (7.5% vs. 1.1%), and *IRS2* (8.4% vs.1.8%) and *GNAS* mutations (6.5% vs. 1.0%). As expected, since *EGFR*-mutant tumors are most often found in never/former light smokers, TMB was significantly lower in the *EGFR*-mutant t-SCLC cohort vs. the *EGFR* WT SCLC cohort (**Fig. 5B**). Similarly, as expected, *EGFR*-mutant t-SCLC tumors were enriched in patients with EAS ancestry (**Fig. 5C**). *EGFR*-mutant t-SCLC cases were also enriched for APOBEC mutational signatures (SBS2 and SBS13) (**Fig. 5D** and **Supplementary Fig. S15)**, indicating that these t-SCLC tumors may present with specific APOBEC-associated mutagenesis, corroborating and expanding on similar analyses in smaller numbers of tumor samples (68). There was no significant difference in mOS between *EGFR*-mutant and wild-type tumors (**Supplementary Fig. S16)**.

**Figure 5:**
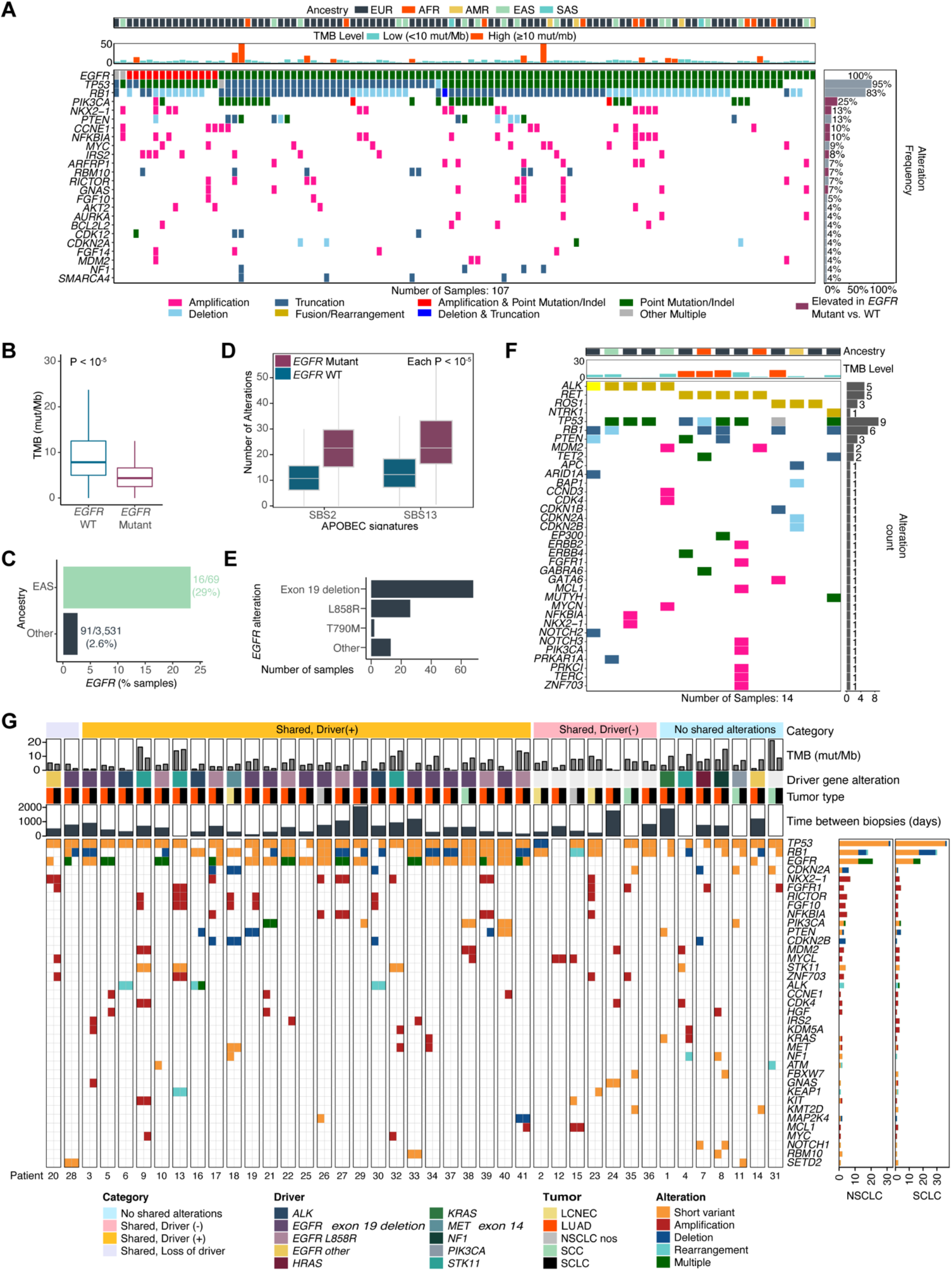
Distinct genetic features of SCLC tumors harboring driver oncogenes traditionally associated with NSCLC. **A**. Oncoplot of the most frequent gene alterations identified in SCLC tumors with kinase-domain mutations in the *EGFR* gene (n=107). Genes are indicated on the left and their alteration frequency on the right. Ancestry, tumor mutation burden (TMB) and microsatellite instability (MSI) for each case, are indicated on top. **B**. Distribution of tumor mutation burden in *EGFR*-mutant and wild-type SCLC tumors. **C**. Comparative prevalence of *EGFR* alterations in patients of East Asian (EAS) ancestry compared to other ancestry groups. **D**. Analysis of the number of mutations representing APOBEC-associated single base substitution signatures (SBS2 and SBS13) in *EGFR* wild-type and mutant SCLC tumors. **E**. Breakdown of the different *EGFR* kinase domain mutations identified in SCLC tumors. **F**. Oncoplot for SCLC tumors with alterations in known oncogenic drivers of NSCLC, excluding *EGFR*. Genes identified to be altered in these cases are shown on the left and their alteration frequency on the right. Tumor mutation burden (TMB) for each case is overlayed on top. **G**. Genomic and clinico-pathological characteristics of 41 patients with paired NSCLC/SCLC samples. Patients were grouped into four categories, based on the patterns of alterations detected: (#1) Shared alterations with Driver (+) in NSCLC and SCLC (orange bar); (#2) Shared alterations with Driver (-) in NSCLC and SCLC (pink bar); (#3) Shared alterations with Driver (+) in NSCLC only and lost/undetected in the matched SCLC (grey bar); (#4) No shared alterations between NSCLC and SCLC (blue bar). Each patient is annotated with the histology, the time between the collection of the NSCLC and SCLC biopsies, the driver alteration (if detected) and tumor mutational burden. Only genes identified to be altered in ≥2 patients are shown. Bar plots showing the total number of patients with each gene alteration is shown on the right, for NSCLC and SCLC samples.

We also observed that our *EGFR*-mutant t-SCLC cohort was enriched for patients with exon 19 deletion (Ex19del, n=68) vs. L858R (n=26) (**Fig. 5E** and **Supplementary Table S17**). Notably, these two “canonical” EGFR mutations occur approximately equal frequency based on numerous previous reports. Indeed, in our dataset as well, Ex19del is detected in 45% of all *EGFR*-mutant NSCLC vs. 64% of all *EGFR*-mutant t-SCLC tumors (**Supplementary Fig. S17**), which can also be seen in a smaller cohort (67), suggesting that tumors with Ex19del may have an enhanced proclivity for lineage transformation.

Interestingly, we also detected other recurrent NSCLC associated oncogenic driver mutations, including *ALK* (n=5), *RET* (n=5), *ROS1* (n=3), and *NTRK1* (n=1) fusions in SCLC tumor samples, suggesting that transformation to SCLC is not exclusive to *EGFR*-mutant NSCLC (**Fig. 5F)**. Co-occurring mutations and TMB were similar to the *EGFR*-mutant t-SCLC cohort.

### Longitudinal analysis of NSCLC to SCLC transformation events reveals potential mechanisms of transformation

41 of the 121 (34%) patients described above had both an NSCLC and an SCLC tumor biopsy genotyped within our dataset. Patterns of shared as well as unique gene alterations were assessed between SNP-matched paired NSCLC and SCLC samples from the same patient across different time intervals, and patients were grouped into potential categories, based on these patterns (**Fig. 5G**). First, there were 7 of 41 (17%) patients in which there were no shared alterations between the NSCLC and SCLC samples. The most likely explanation is that these tumors developed independently within the patients (**Fig. 5G**, light blue box). Second, there were 7 of 41 patients (17%) in which the NSCLC and SCLC samples shared alterations, but no previously described NSCLC driver gene alteration was detected in either biopsy (**Fig. 5G**, pink box). These patients would not have been treated with oncogene targeted therapies, suggesting that other treatment modalities, such as chemotherapy and immunotherapy, can also drive SCLC transformation. Third, in 25 of 41 patients (61%), the NSCLC and SCLC tumor samples both contained a driver mutation previously associated with NSCLC (**Fig. 5G**, orange box). This was the largest cohort within the matched, paired patient NSCLC-SCLC samples, and our results are concordant with previous data showing that cases of t-SCLC typically retain the original NSCLC driver mutation (24,67,68). Finally, there were 2 patients for which a driver mutation was detected in the NSCLC sample but not in the SCLC sample (**Fig. 5G**, grey box). In both cases, the NSCLC and SCLC samples shared other alterations, suggesting that the driver mutation was lost during the transformation process, for example by recombination with the wild-type allele.

Taken together, our analysis of >100 putative t-SCLC reveals that lineage plasticity may arise from multiple different molecular cohorts of NSCLC, including *EGFR*-mutant and fusion kinase positive NSCLC. Furthermore, analysis of paired samples demonstrates that putative SCLC transformation may occur at variable lengths of time from the original NSCLC diagnosis and suggests that time on therapy may not be a factor for stratifying which patients may have t-SCLC at the time of disease progression. With an increased use of inhibitors targeting these oncogenic drivers in the clinic, it is likely that more cases of t-SCLC coming from these other types of NSCLC will be identified in the future and a broader implementation of rebiopsy at the time of acquired therapeutic resistance may prove important.

## DISCUSSION

Herein, we present a genomic analysis of the largest cohort of SCLC tumors evaluated to date, encompassing 3,600 tumor specimens. This study is unique in many ways: (a) The SCLC samples in our cohort have been predominantly obtained from community sites throughout the United States, representing a more typical “real-world” cohort of SCLC. (b) Genomic data are tied to clinical data (including survival) and predicted ancestry data, parameters that have been limited in most previous SCLC studies. This allowed us for the first time to investigate genetic differences in SCLC tumors based on ancestry. (c) To the best of our knowledge, our study contains the analysis of the largest number of SCLC tumors from African ancestry to date (n=256, 7.1% of the entire cohort). (d) As a result of the large sample size, we were able to evaluate mutational status by anatomical location of the tumor and show site specific enrichment of certain genomic alterations. (e) This study contains the largest cohort studied of putative “transformed SCLC” (t-SCLC) (n=121, including 107 *EGFR*-mutant cases).

A major limitation of our study is the number of genes for which alterations is queried, with deep sequencing of exons from up to 324 cancer-related genes and select introns from up to 34 genes frequently rearranged in cancer. Because SCLC is a cancer type with unique biology, it is possible that some frequently altered genes are not included in the list of genes analyzed. For example, whole-genome sequencing identified recurrent loss-of-function alterations in genes coding for the RB family members p107 and p130 (7), but these two genes (*RBL1* and *RBL2*) are not included in our panel.

Another limitation of our data is the lack of RNA/protein expression profiles for the tumors analyzed. It is possible that some gene alterations identified at the DNA level do not result in changes in gene/protein expression in SCLC cells, including some of the new rearrangements identified. Despite these limitations, we have made several key observations related to SCLC pathobiology which were not possible from previous studies due to sample size, including new genetic subgroups, site-specific mutational patterns, and insights into histological transformation (summarized in **Fig. 6**).

**Figure 6:**
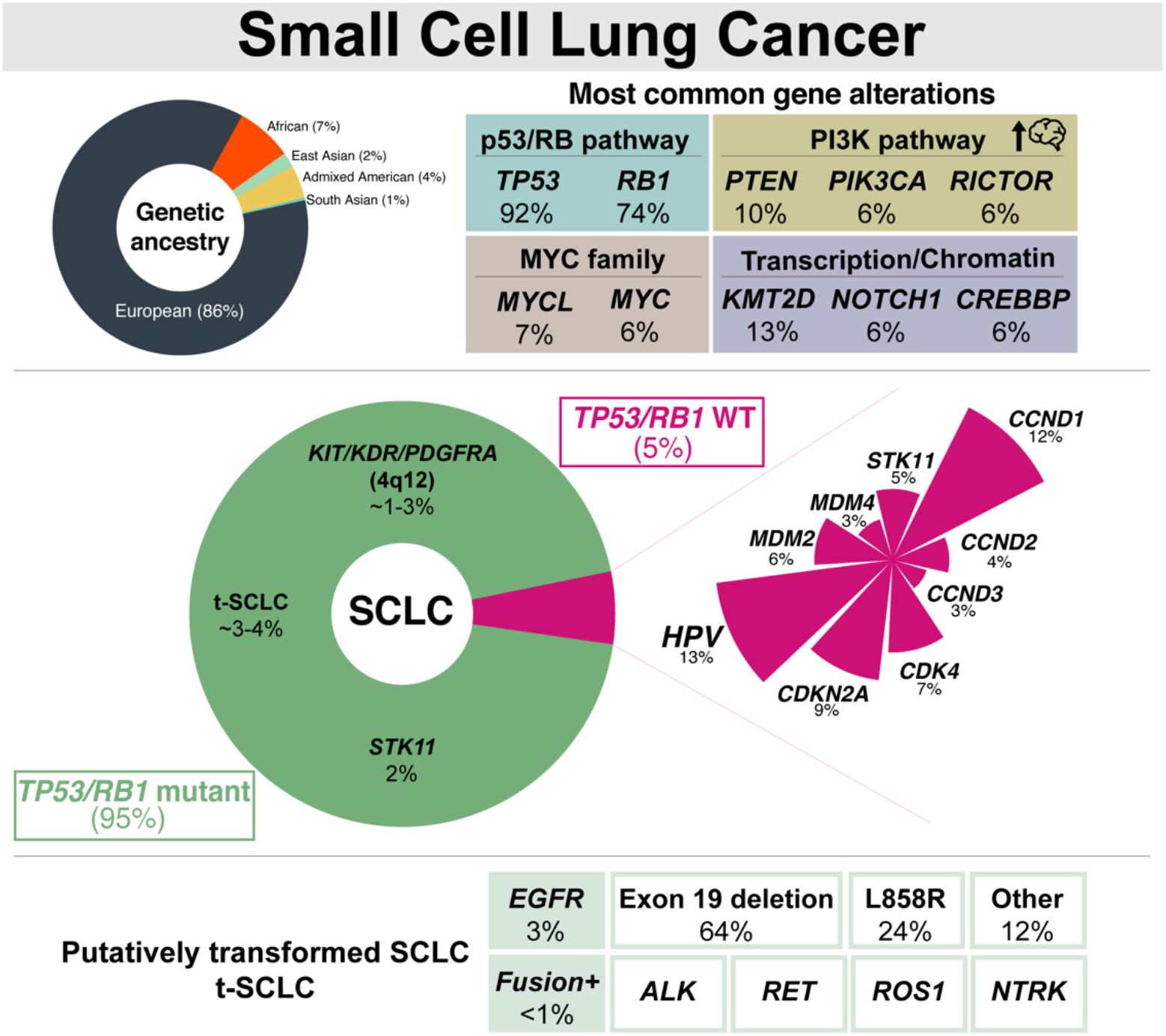
Overview and summary of the main findings from this analysis of 3600 cases of SCLC. Schematic representation highlighting the main findings from our integrative analysis of 3600 patients with SCLC with diverse genetic ancestry.

Our work underscores the existence of genetic subtypes in SCLC and key genetic determinants of patient outcomes. Some of these subtypes may be rare or very rare, but patients with these tumors may greatly benefit from personalized treatments. For example, HPV-positive *TP53*/*RB1* wild-type tumors may represent <0.1% of cases, but these tumors may be uniquely responsive to strategies targeting the virus or re-activating p53 function to induce cell death. As another example, *STK11* mutations were detected in 1.7% of our entire cohort, were enriched in patients of AFR ancestry, and were associated with a decreased OS compared with the *STK11* wild-type cohort. Numerous studies have now shown that *STK11* mutations are associated with decreased response to ICI therapy in NSCLC (62-64), and it is possible that the same is true in SCLC.

Therapeutic selection based on tumor mutational status has revolutionized the care of patients with NSCLC, leading to the development of numerous new drugs and implementation of personalized treatment approaches which have become the accepted standard of care for patients with NSCLC around the world. However, implementation of personalized therapies for patients with SCLC has remained elusive. Classification of SCLC into subtypes defined by transcriptional programs has been postulated, with two main subtypes emerging - neuroendocrine (NE) and non-neuroendocrine (non-NE) (50). However, the prognostic, predictive, and clinical significance of these subtypes is not well defined. Furthermore, how specific mutations influence the subtype landscape remains unclear. NE tumors are enriched for alterations in *RB1, NOTCH1, MYCL1*, and chromatin modifier genes and these NE tumors may have improved responses to replication stress inhibitors and poorer responses to ICI therapy compared to non-NE tumors (14). However, in this same study, there were no differences in overall survival between patients with NE vs. non-NE tumors, suggesting that the transcriptomic type alone may be insufficient for stratification.

Importantly, our large “real-world” cohort, while composed of a majority of patients from European ancestry, is also the most diverse cohort of SCLC tumors analyzed to date, which allowed us for the first time to investigate genetic differences in SCLC tumors based on ancestry. For instance, we previously discussed the need to better investigate genetic, environmental, and socioeconomic factors associated with SCLC development in various populations, including in Black patients with SCLC (69). SCLC tumor samples harboring activating mutations within the EGFR kinase domain were found in all ancestry groups but enriched in patients of East Asian ancestry, as expected (70, 71). More surprisingly, when we analyzed 256 SCLC tumors in patients of African ancestry, we found that these tumors were more likely to be wild-type for *RB1* and *TP53*, and mutant for *STK11*, suggesting that populations of different ancestry may be enriched for different genetic subtypes of SCLC. These observations may help guide genetic testing and clinical decisions. The challenges associated with capturing and analyzing self-reported race in the clinical trial setting has been well-described in the literature (72). The use of genomic ancestry in this real-world population offers a model for describing genomic differences across ancestral populations and contributes to a growing number of similar studies in other cancer types (73-75).

Our cohort of patients with matched NSCLC and SCLC samples also extends on previous observations related to lineage plasticity in several ways. Notably, these data show that SCLC transformation can occur across multiple different NSCLC driver mutations, not just *EGFR*-mutant NSCLC. This is important because tumor re-biopsy at the time of disease progression is not currently mandated for all molecular cohorts of NSCLC. Yet, at present, tumor biopsy is the only way to definitively diagnose SCLC transformation, and a finding of SCLC in a patient previously diagnosed with NSCLC would change clinical management in terms of therapeutic selection. Current clinical trials attempting to prevent SCLC transformation by combining targeted therapy with platinum-etoposide chemotherapy (traditionally used for SCLC) necessitate that the NSCLC tumor harbors both *TP53* and *RB1* mutations (NCT03567642). This is important based on our data that putative transformation from NSCLC to SCLC does not strictly require both mutation in *TP53* and mutation in *RB1*.

Overall, these data are expected to serve as a catalyst for additional laboratory-based and clinical research studies that will further our knowledge of SCLC biology and the biology of other types of neuroendocrine tumors as well as provide much needed advances in treatment paradigms for patients with SCLC.

## METHODS

### Targeted next generation sequencing (NGS) of SCLC tumors

Formalin-fixed, paraffin embedded (FFPE) tissue sections of SCLC from 3,600 patients were profiled using FoundationOne^®^ (N=1,515) or FoundationOne^®^CDx (N=2,085) comprehensive genomic profiling (CGP) assays in a Clinical Laboratory Improvement Amendments (CLIA)-certified, College of American Pathologists (CAP)-accredited laboratory (Foundation Medicine Inc., Cambridge, MA, USA). Briefly, a minimum of 50 ng of DNA was extracted from FFPE sections and CGP was performed on hybridization-captured, adaptor ligation-based libraries to a median exon coverage depth of > 500X for exons of up to 324 cancer-related genes and select introns from 34 genes frequently rearranged in cancer (**Supplementary Table S2**).

A multi-stage pathology review was performed prior to and after sequencing. Prior to sequencing, board-certified pathologists on staff at Foundation Medicine reviewed the submitted pathologic diagnosis of each case and examined hematoxylin and eosin (H&E)-stained slides. Tumor type assignment for each case was performed based on the submitting diagnosis and re-review of the H&E. If necessary, pathologist-directed macro-dissection to achieve >20% estimated percent tumor nuclei (%TN) in each case was performed, where %TN is defined as 100 times the number of tumor cells divided by total number of nucleated cells. Approval for this study, including a waiver of informed consent and HIPAA waiver of authorization, was obtained from the Western Institutional Review Board (Protocol No. 20152817).

### Identification of genomic alterations

Different classes of genomic alterations, comprising short variants (base substitutions and small insertions/deletions), copy number alterations (focal amplifications and homozygous deletions), and rearrangements were identified, using the approach described previously (76). Tumor mutational burden (TMB) was determined on 0.8–1.2 Mb of sequenced region and samples with a TMB of at least 10 mutations/Mb were classified as TMB-high (77).

### Detection of chromosomal arm-level aneuploidy

A copy number modeling algorithm was used to identify chromosome-arm level aneuploidy. Briefly, for each sample, the algorithm utilizes the coverage profile for regions of the genome targeted in the assay, normalized to a process-matched normal control, to model the copy number of each segment. The minor allele frequencies of up to 59,622 single nucleotide polymorphisms (SNPs) distributed across each segment were used, along with the normalized intensities to identify regions under aneuploidy (gain, loss). If over 50% of a chromosome arm exhibited a gain or a loss, it was classified as a chromosomal arm-level aneuploidy event. Additionally, the chromosome arm score (Charm) for tumor suppressors (CharmTSG) and oncogenes (CharmOG) described by Davoli and colleagues (29), was used to perform correlation analyses against the identified chromosomal arm-level aneuploidy events identified in our cohort.

### Prediction of patient genomic ancestry

Ancestry for each patient was predicted using single nucleotide polymorphisms (SNP) from the targeted NGS assay that overlapped with those captured in the Phase 3 1000 Genomes (78). This SNP-based approach was projected down to five principal components, that were used to train a random forest classifier to identify the following ancestry groups: European (EUR), African (AFR), East Asian (EAS), South Asian (SAS) and admixed American (AMR), as described previously (79).

### Analysis of genomic and clinical patterns in specific subpopulations

Differences in prevalence of gene alterations and biomarkers were assessed using a Fisher’s exact test with false discovery rate (FDR)-based correction for multiple testing. Only genes that were targeted on both assay platforms were assessed (**Supplementary Table S2**). Additionally, for continuous variable biomarkers (e.g., TMB), Wilcoxon rank sum test was used to assess the differences between different subgroups.

### Identification of mutational signatures

We utilized a pooled approach to assess mutational signatures in different SCLC alteration subgroups from the targeted panel assay. Comparisons were performed between *EGFR*-mutant and *EGFR*-WT as well as *TP53/RB1*-mutant and *TP53/RB1*-WT samples. Mutagenic signatures are typically considered to be additive and therefore, difference between the two groups was used to understand the additional effect of mutations. All variants that were predicted non-germline and variants of unknown significance were pooled for this analysis. The contributions of the known COSMIC v3.2 signatures were obtained as described previously (80, 81). The number and percentage of mutations attributed to each mutational signature was calculated relative to the total number of pooled mutations. The stability of this pooled approach was evaluated by resampling the cohorts to a jackknife sample size, defined as the mutations from 30 samples. Both cohorts were resampled without replacement 1,000 times and signature attribution was assessed on each of the resampled pools of mutations. This provides both the median and 95% confidence interval for the contribution of all known COSMIC v3.2 mutation signatures in each cohort. Samples with a very high tumor mutational burden (≥ 50 mutations/Megabase) were excluded from this pooled mutation signature analysis.

### Analysis of clinical outcomes

This study used the nationwide (US-based) de-identified Flatiron Health (FH)-Foundation Medicine (FMI) SCLC clinico-genomic database (CGDB). The de-identified data originated from approximately 280 United States cancer clinics (∼800 sites of care). Retrospective longitudinal clinical data, derived from electronic health records comprising patient-level structured and unstructured data, were curated via technology-enabled abstraction, and linked to the genomic data obtained from the CGP test at FMI using de-identified, deterministic matching(82). This study included 678 patients with a SCLC diagnosis who received care in the FH network between 01/2011 and 09/2021 and underwent tissue biopsy-based CGP (FoundationOne^®^ or FoundationOne^®^CDx) between 10/2012-9/2021. Institutional Review Board approval of the study was obtained prior to study conduct and included a waiver of informed consent.

Overall survival as calculated from the date of SCLC diagnosis to either date of death or date of last clinic visit. Patients were treated as at risk of death only after the later of their first sequencing report date and their second visit in the Flatiron Health network on or after January 1, 2011, as both are requirements for inclusion in the database. Treatment start dates were determined by oncologist-defined, rule-based lines of therapy. Time to next treatment was calculated from the first line treatment start date to either the second line treatment start date or date of death if death occurred prior to the start of second line treatment. Patients without an event were censored at the date of their last clinic visit. All time to event outcomes were assessed by log-rank test and univariate Cox proportional hazards models. Analyses were performed on R software version 4.0.3.

### Viral detection

A *de novo* assembly of off-target sequencing reads left unmapped to the human reference genome (hg19) was performed as described previously (59). These assembled contigs were competitively aligned by BLASTn to the NCBI database of viral nucleotide sequences to detect oncoviruses, including the human papillomavirus (HPV) types and Merkel cell polyomavirus. Contigs ≥80 nucleotides in length and with ≥97% identity to the BLAST sequence were determined to be positive for viral status.

### Investigation of liquid biopsies

A total of 81 SCLC cases profiled on the FoundationOne^®^Liquid CDx assay were examined for their mutational patterns (83). Quantification of the ctDNA fraction was performed using two complementary methods: a tumor fraction estimator (TFE) and the maximum somatic allele frequency (MSAF) method (84). TFE is based on a measure of tumor aneuploidy and MSAF uses allele fraction from somatic coding alterations to estimate ctDNA fraction. Patterns of ctDNA fraction in this SCLC cohort were contrasted to 4,573 non-small cell lung cancers profiled on the same assay.

### Funding

This work was supported by the National Institutes of Health grant numbers CA231851, CA217450, and CA231997 to J.S. and CA217450, CA224276, and CA233259 to C.M.L. C.M.L. was also partially supported by a LUNGevity Foundation award with funds raised by EGFR Resisters for the 2021 EGFR Resisters/LUNGevity Foundation Lung Cancer Research Award Program.

### Author contributions

S.S., J.S., and C.M.L. designed the study and interpreted the results. S.S., J.A.M., M.M., R.S., D.L, Z.F., E.E., and J.N performed data analyses and curation. J.M., P.S.H and G.M.F provided supervision and writing review and editing. S.S. and M.M. prepared the figures. S.S., J.S., and C.M.L. wrote the manuscript with contributions from all authors.

## Supporting information

Sivakumar et al_Supplemental Tables

## Acknowledgements

The authors thank members of the Lovly and Sage labs, the Foundation Medicine scientific review team, and members of the U54 SCLC Consortia for critical comments on the manuscript.

## SUPPLEMENTARY FIGURES

**Figure S1:**
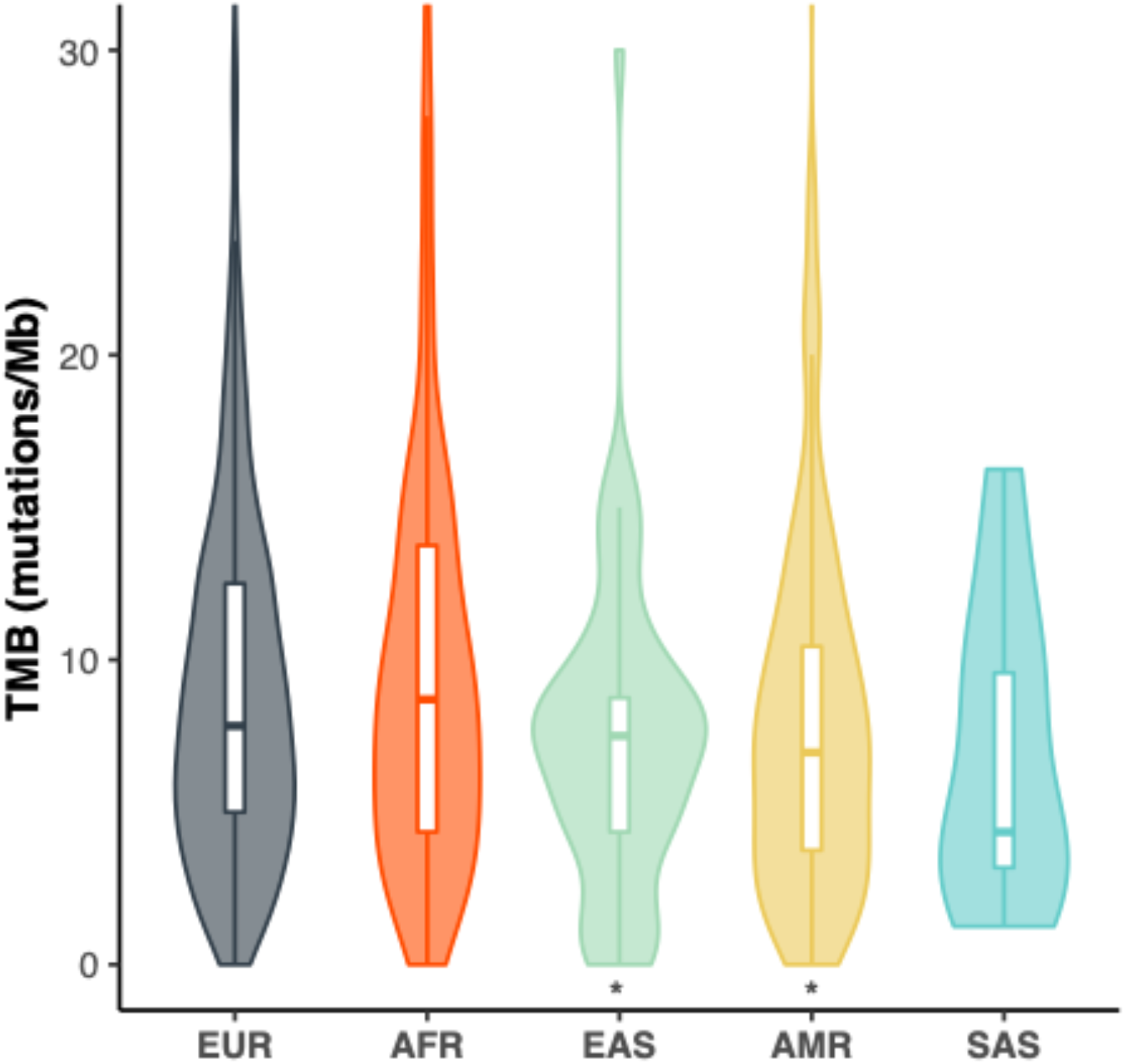
Patterns of tumor mutational burden across different ancestry groups. Violin plots showing the distribution of tumor mutation burden (TMB) in SCLC tumors across various ancestry groups. EUR, European; AFR, African; EAS, East Asian; AMR: Ad-mixed American; SAS, South Asian. Comparison of the TMB distribution between each ancestry group was performed against the EUR group using a Wilcoxon test (p value threshold: *: 0.05).

**Figure S2:**
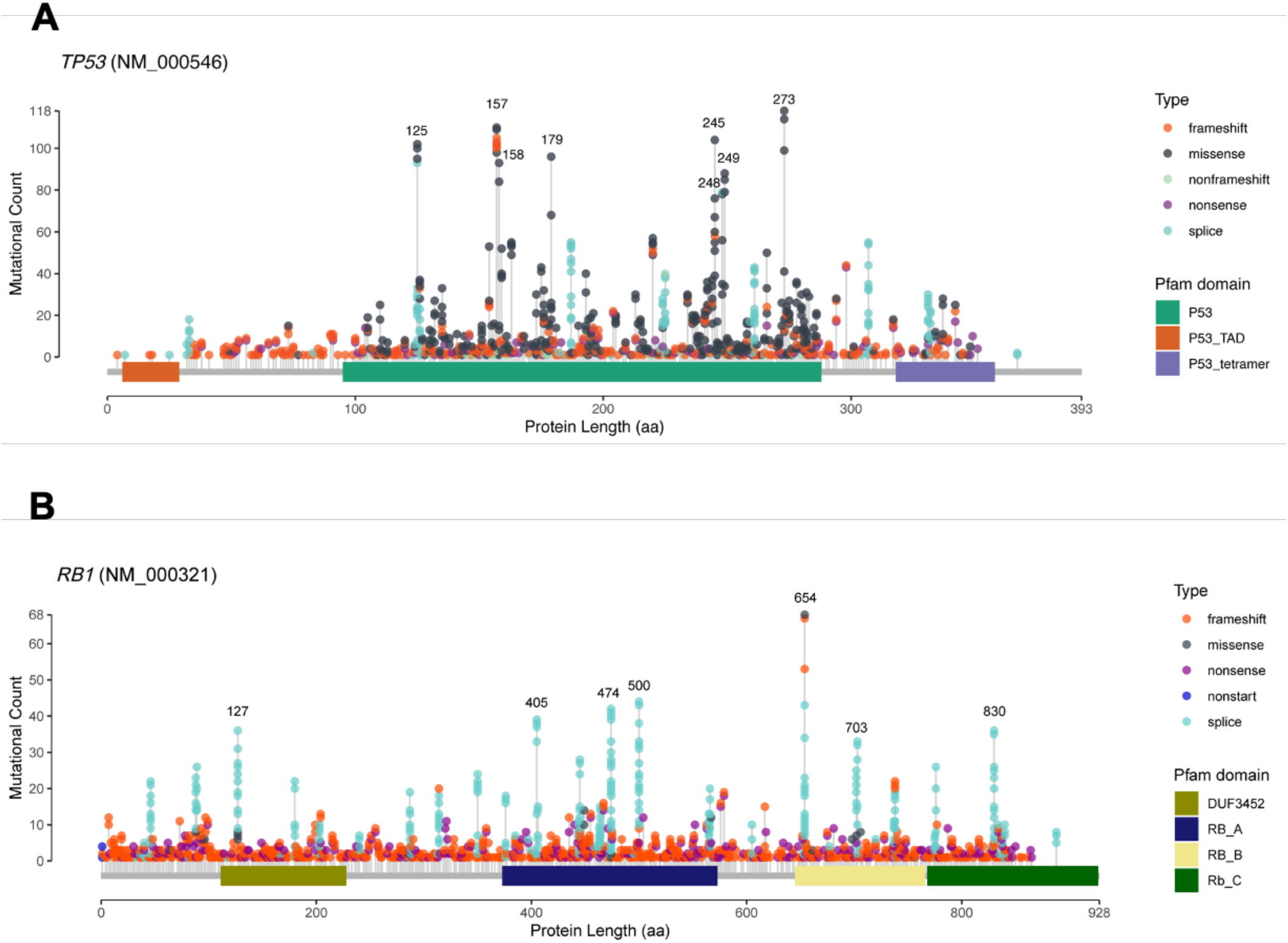
Spectrum of *TP53* and *RB1* mutations in SCLC. Lollipop plot representation of all identified (**A**) *TP53* and (**B**) *RB1* mutations, including single nucleotide variants, short insertions/deletions as well as splice alterations, in SCLC tumors across the protein length. The most frequently mutated regions/codons are labelled in each plot.

**Figure S3:**
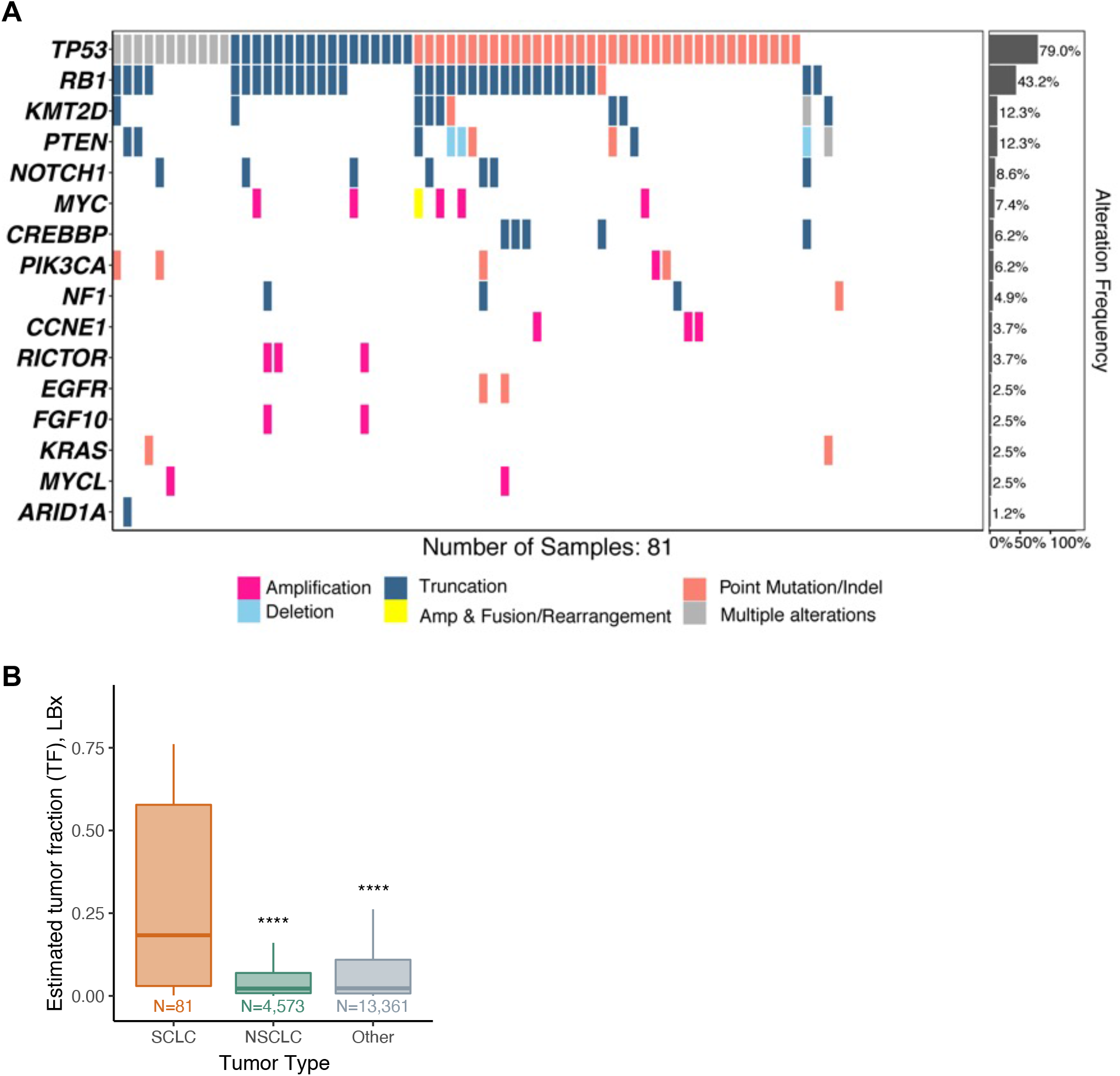
Gene alteration landscape detected from targeted sequencing of liquid biopsies. **A**. Frequent alterations identified in SCLC liquid biopsies (N=81). Genes are indicated on the left and their alteration frequency on the right. **B**. Boxplots showing the estimated tumor fraction (see Methods) for SCLC, NSCLC, and other tumors profiled using liquid biopsies. Comparison of the distribution of estimated tumor fraction between SCLC tumors and other tumor groups was performed using a Wilcoxon test (P value thresholds: *: 0.05, **: 0.01, ***: 0.001, ****: 0.0001).

**Figure S4:**
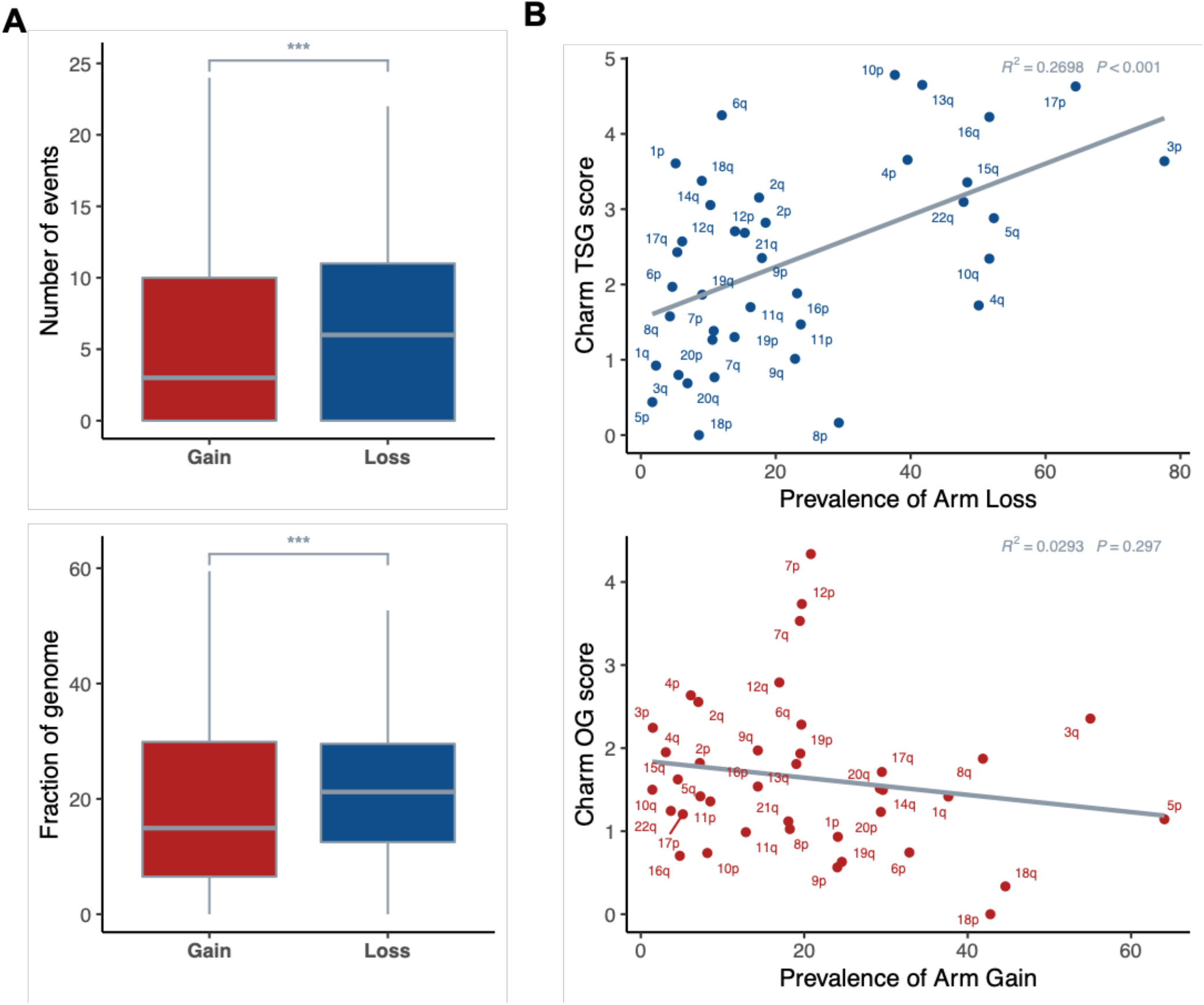
Chromosomal loss events are frequent in SCLC and are preferentially enriched for tumor suppressor genes. **A**. Boxplots showing the burden of chromosome arm-level gain and loss events based on number of events (top) and fraction of the genome (bottom). **B**. Prevalence of chromosome-arm level loss and gain events compared to the Charm tumor suppressor (TSG) and oncogene (OG) scores for each arm.

**Figure S5:**
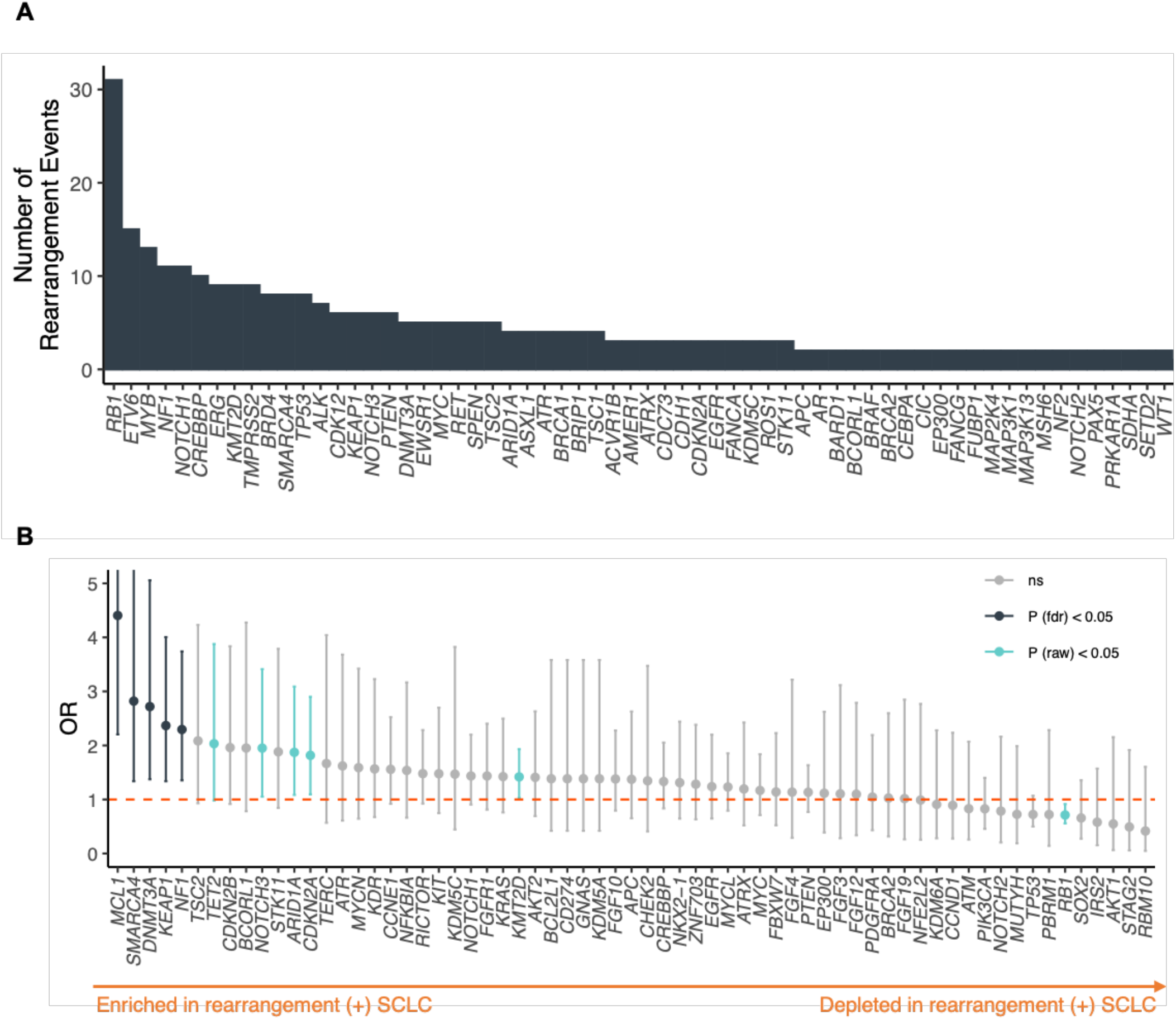
SCLC tumors exhibit recurrent gene rearrangements with therapeutic potential. **A**. Number of recurrent gene rearrangement events in the overall cohort of SCLC tumors. Genes are ordered based on the number of detected rearrangements. **B**. Comparison of prevalence of gene alterations in SCLC tumors with gene rearrangement events against SCLC tumors without any identified gene rearrangement events. Genes are ordered by their odd’s ratio (OR). Comparison of gene alteration prevalence between the groups was performed using a Fisher’s exact test with FDR-based correction (ns: not statistically significant).

**Figure S6:**
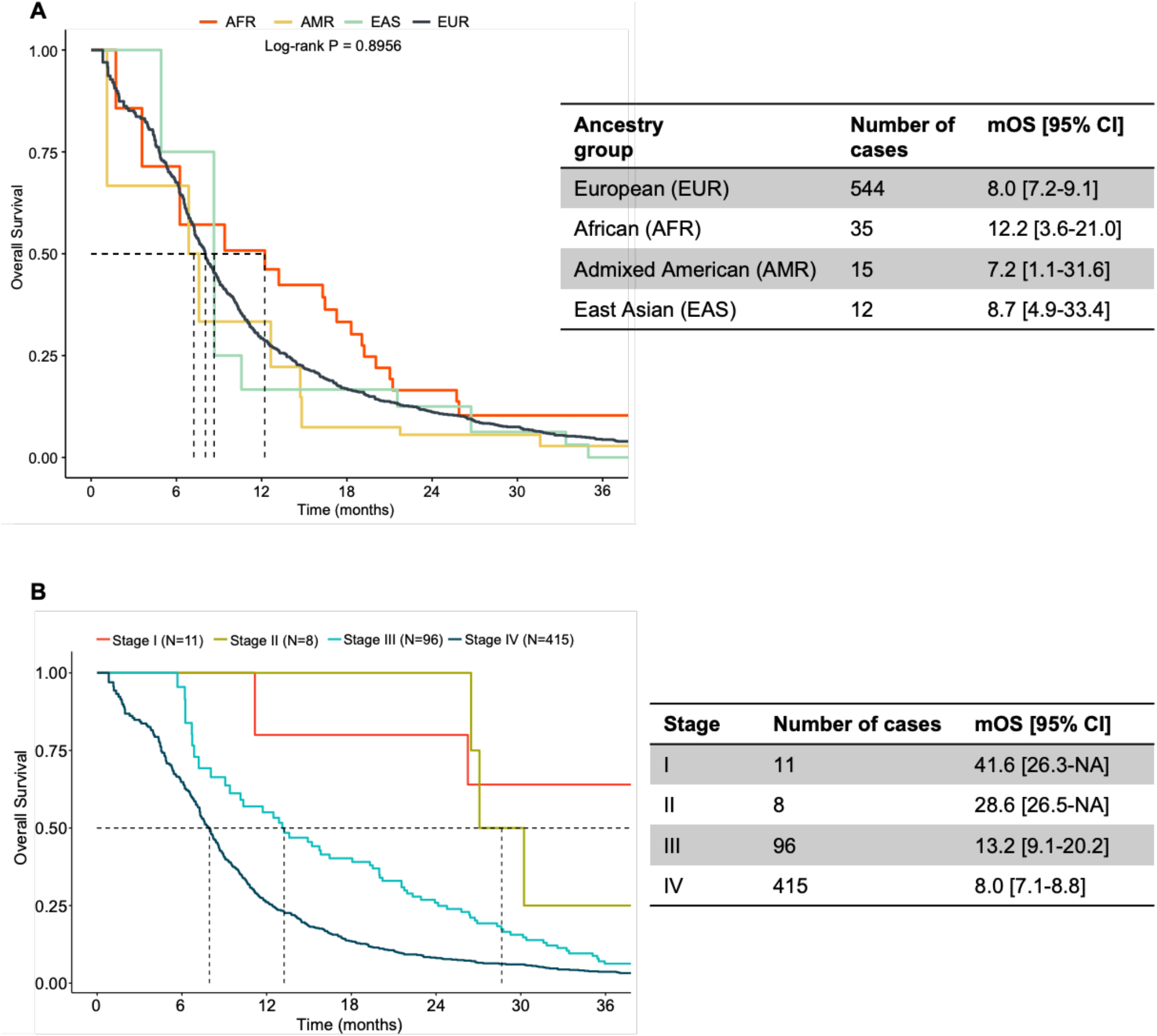
Trends of overall survival across different ancestry groups and stage in the clinical cohort. **A.** Kaplan-Meier plot displaying the overall survival across ancestry groups in the SCLC clinical cohort. EUR, European; AFR, African; EAS, East Asian; AMR: Ad-mixed American. South Asian ancestry cohort was excluded due to low counts. **B.** Kaplan-Meier plot displaying the overall survival of patients with SCLC diagnosed at different stages of tumor progression in the clinical cohort. Patients with unknown stage information (N=77) were not plotted.

**Figure S7:**
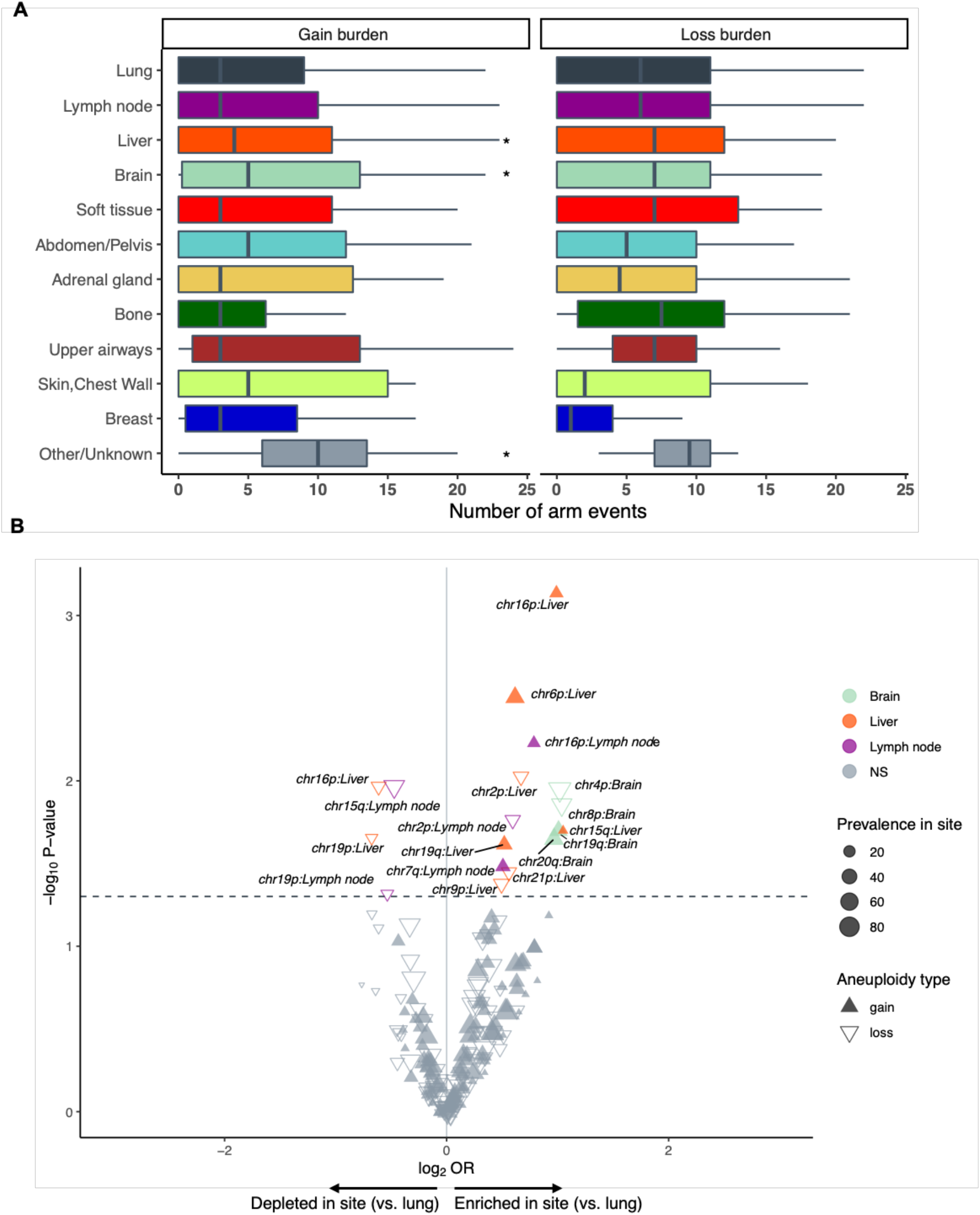
Liver and Brain SCLC metastases are enriched for chromosomal arm-level gains compared to lung-biopsied SCLC tumors. **A**. Burden of chromosomal arm-level events, estimated as the number of chromosome arms with a copy number event, in lung-biopsied and other metastatic site-biopsied SCLC tumors. **B**. Volcano plot displaying the comparative prevalence of chromosomal arm-level changes across the most frequent metastatic sites (brain, lymph node, liver) compared to lung-biopsied SCLC tumors. These comparisons were not corrected for multiple hypothesis testing.

**Figure S8:**
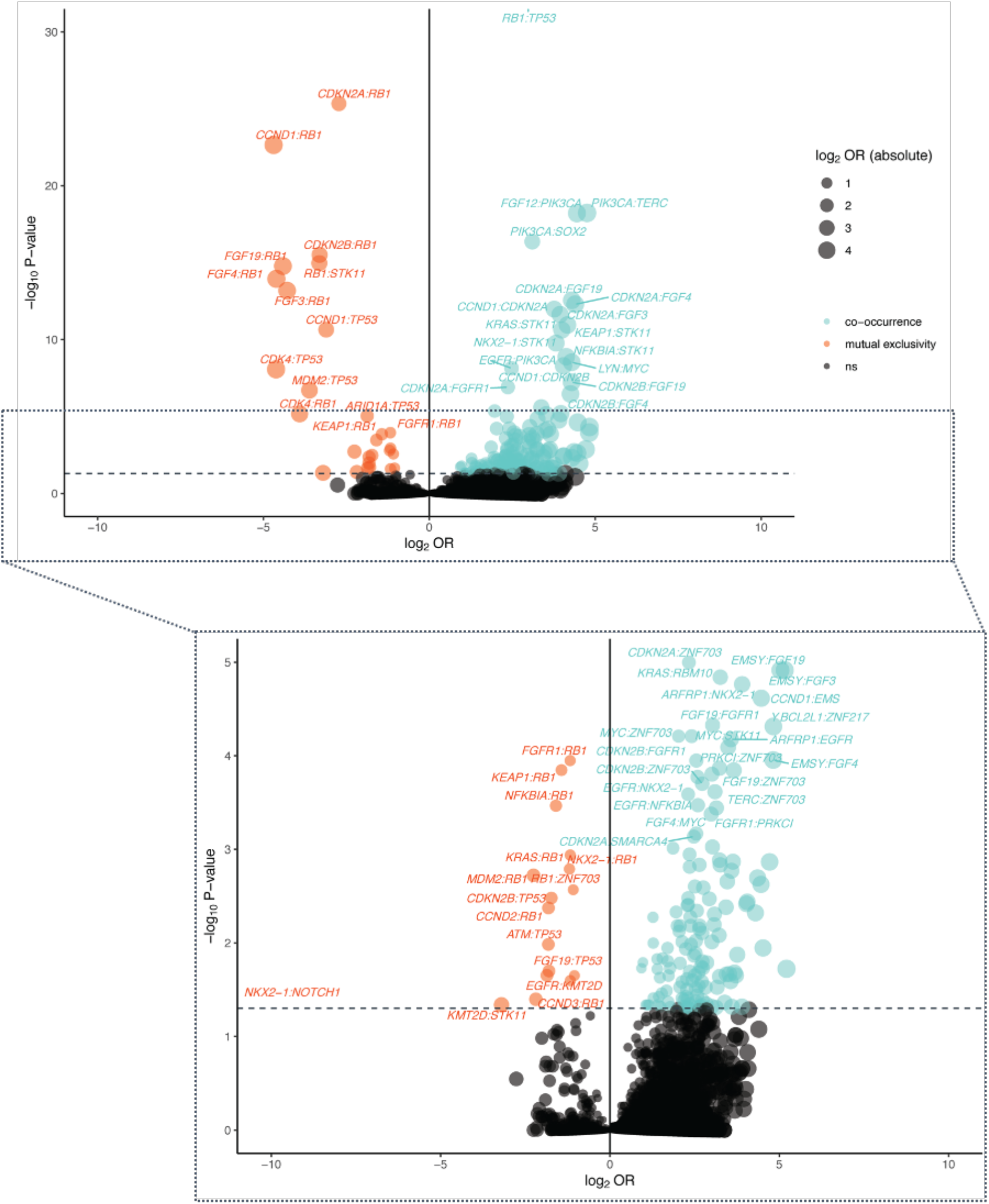
SCLC tumors present with unique co-occurring and mutual exclusive patterns of gene alterations. Volcano plots from the analysis of co-occurrence (blue) and mutual exclusivity (orange) between gene alterations in SCLC tumors. Each point represents a pair of genes.

**Figure S9:**
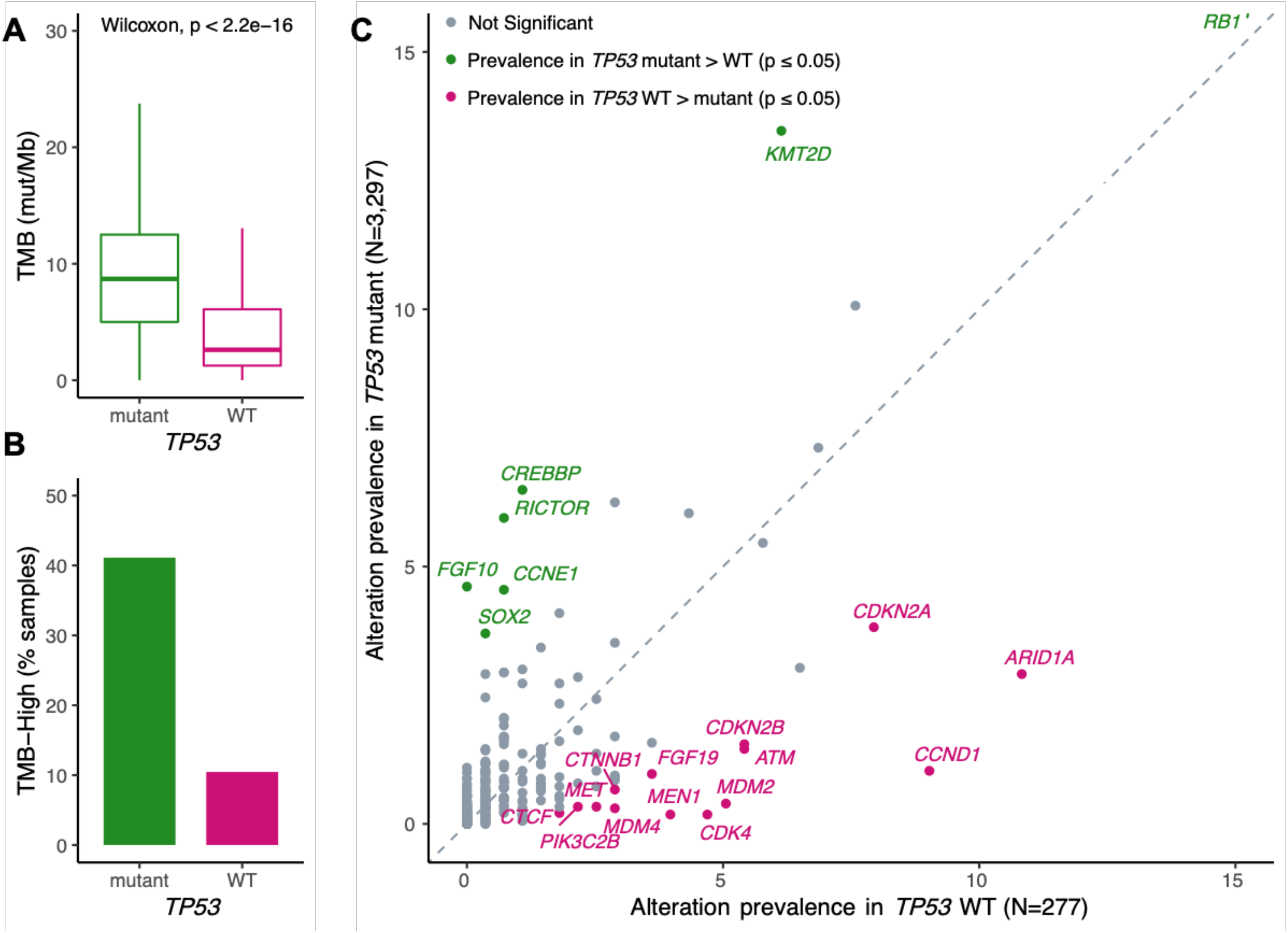
SCLC tumors with wild-type *TP53* have a lower tumor mutational burden and present with specific gene alterations. **A**. Distribution of tumor mutation burden (TMB, mutations/Mb) in SCLC tumors wild-type and mutant for *TP53*. **B**. Prevalence of TMB-high status (≥10 mutations/Mb) in SCLC tumors wild-type and mutant for *TP53*.**C**. Comparison of gene alteration prevalence in *TP53*-mutant and wild-type SCLC tumors.

**Figure S10:**
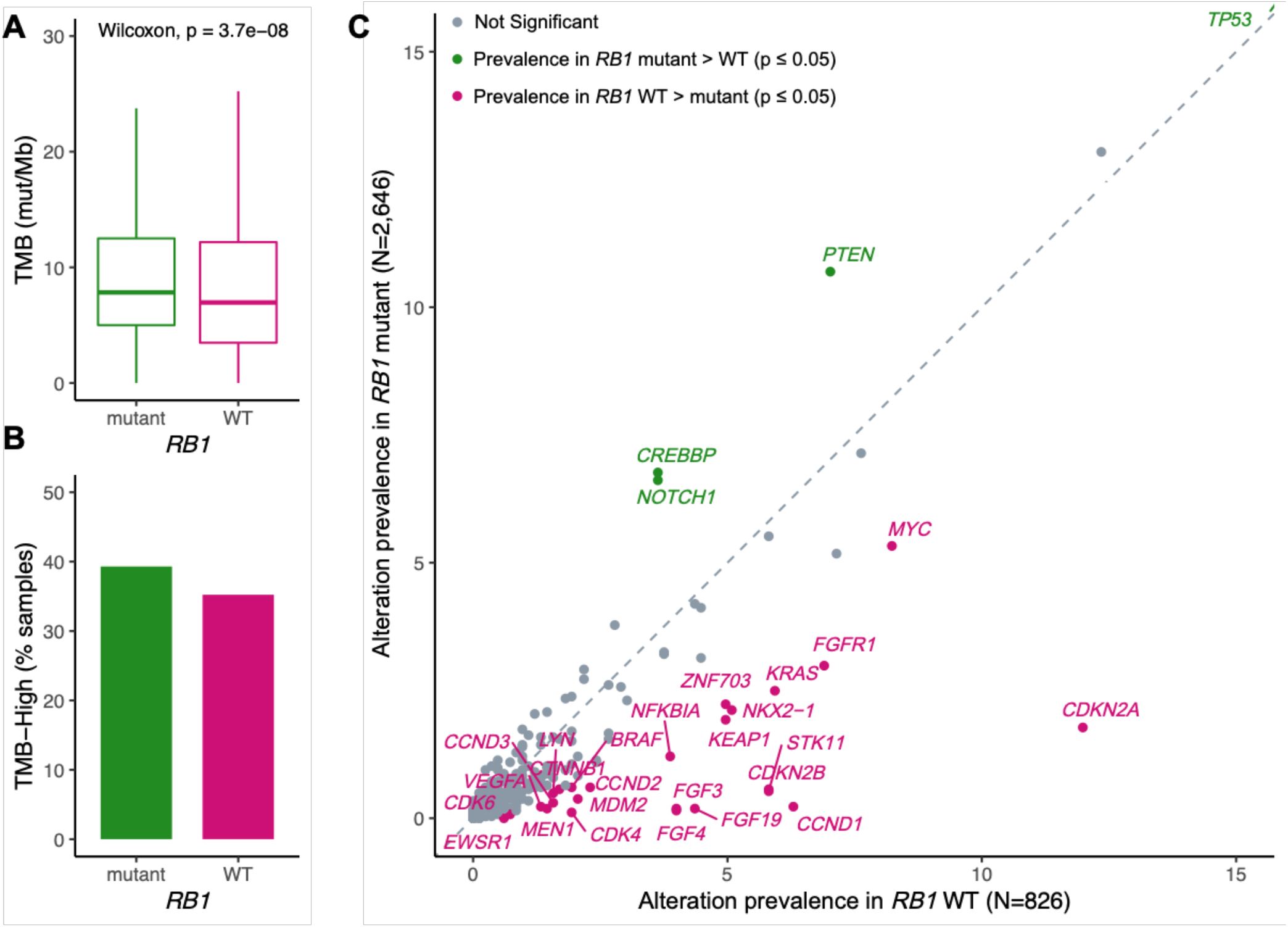
SCLC tumors with wild-type *RB1* have a lower tumor mutational burden and present with specific gene alterations. **A**. Distribution of tumor mutation burden (TMB, mutations/Mb) in SCLC tumors wild-type and mutant for *RB1*. **B**. Prevalence of TMB-high status (≥10 mutations/Mb) in SCLC tumors wild-type and mutant for *RB1*. **C**. Comparison of gene alteration prevalence in *RB1*-mutant and wild-type SCLC tumors.

**Figure S11:**
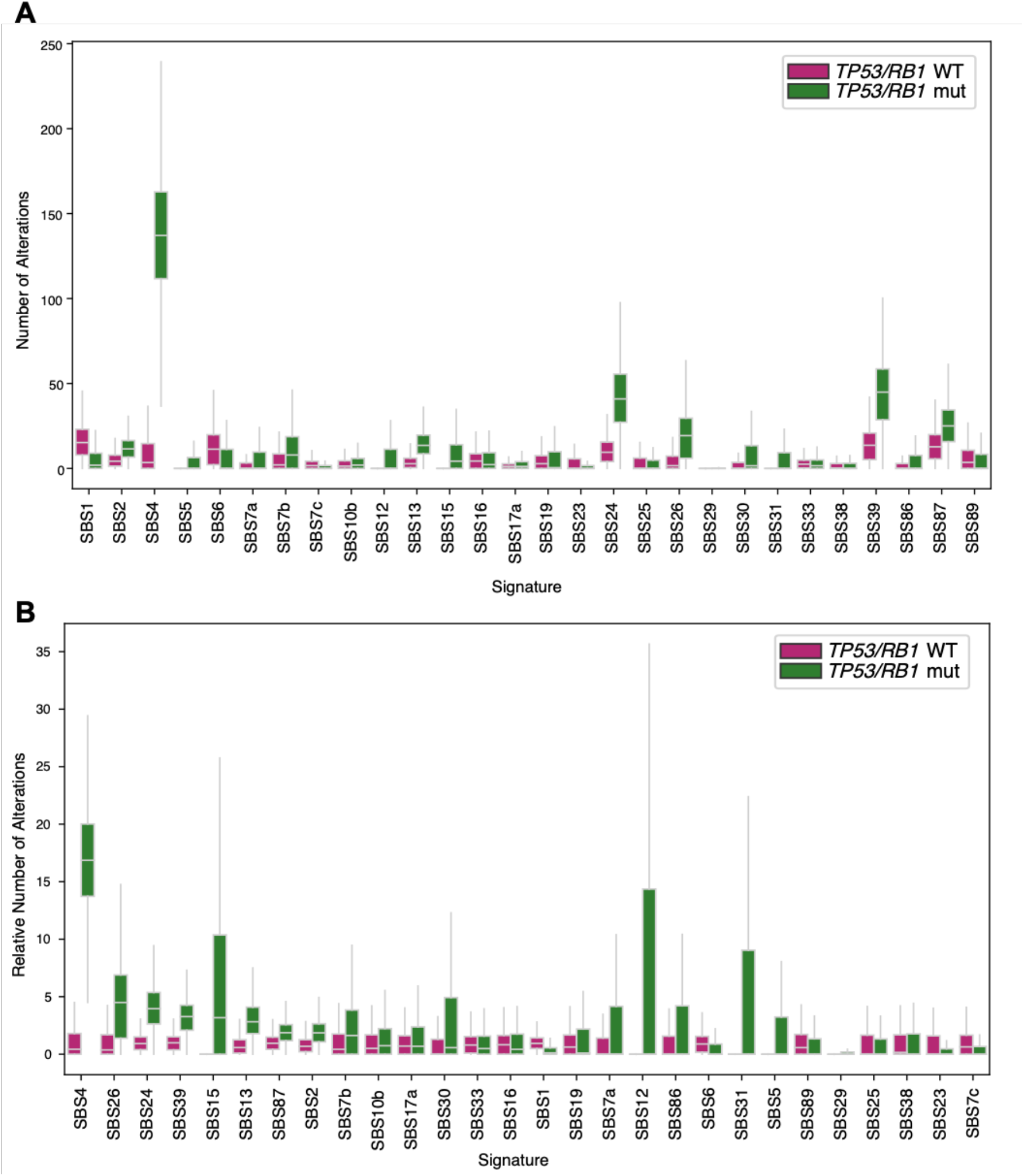
A smoking-associated mutational signature is rare in *TP53/RB1* WT SCLC tumors. Analysis of the absolute number of alterations (**A**) and relative number of alterations (**B**) attributable to different single base substitution (SBS) signatures in *TP53/RB1* wild-type and mutant tumors. Tobacco-associated SBS4 signature is rare among *TP53/RB1* wild-type tumors, in comparison to *TP53/RB1* mutant tumors. The COSMIC annotations for the displayed mutational signatures include: SBS1 (clock-like), SBS2 (APOBEC), SBS4 (tobacco smoking), SBS5 (clock-like), SBS6 (DNA mismatch repair deficiency), SBS7a, SBS7b, SBS7c (ultraviolet light exposure), SBS10b (polymerase deficiency), SBS12 (unknown), SBS13 (APOBEC), SBS15 (DNA mismatch repair deficiency), SBS16, SBS17a, SBS19, SBS23 (unknown), SBS24 (aflatoxin exposure), SBS25 (chemotherapy treatment), SBS26 (DNA mismatch repair deficiency), SBS29 (tobacco chewing), SBS30 (DNA base excision repair deficiency), SBS31 (platinum chemotherapy treatment), SBS33 (unknown), SBS38 (indirect effect of ultraviolet light), SBS39 (unknown), SBS86 (unknown chemotherapy treatment), SBS87 (thiopurine chemotherapy treatment), SBS89 (unknown).

**Figure S12:**
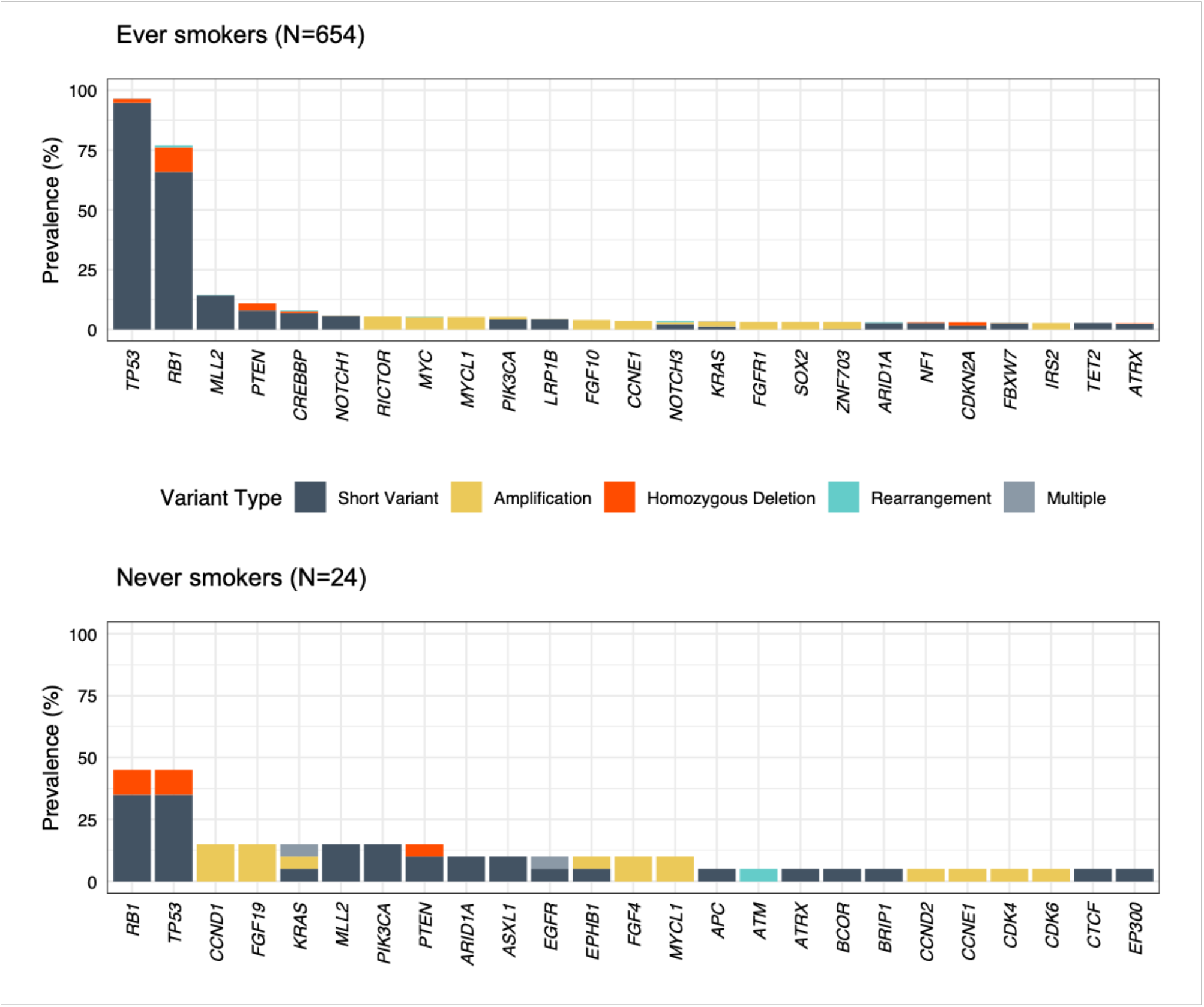
T*P*53 and *RB1* alterations are less frequent in never smokers. Prevalence of mutations in recurrently mutated genes, in ever smokers and never smokers from the clinical cohort.

**Figure S13.**
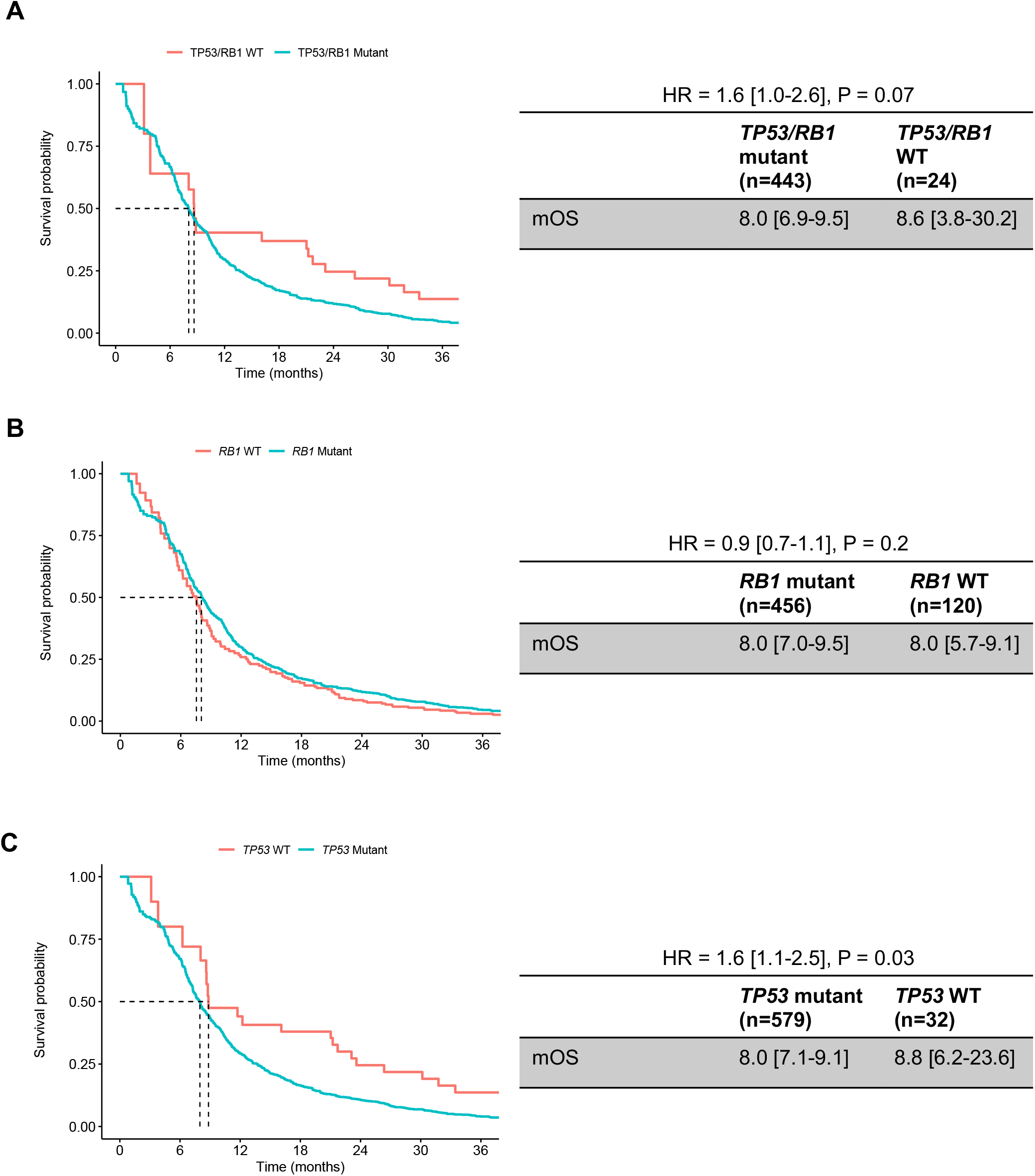
Patterns of overall survival based on *TP53* and *RB1* alteration status. Survival probability in SCLC tumors that are wild-type or mutant for (**A**) *TP53* and *RB1* (**B**) *RB1* (**C**) *TP53*. HR, hazard ratio; mOS, median overall survival.

**Figure S14.**
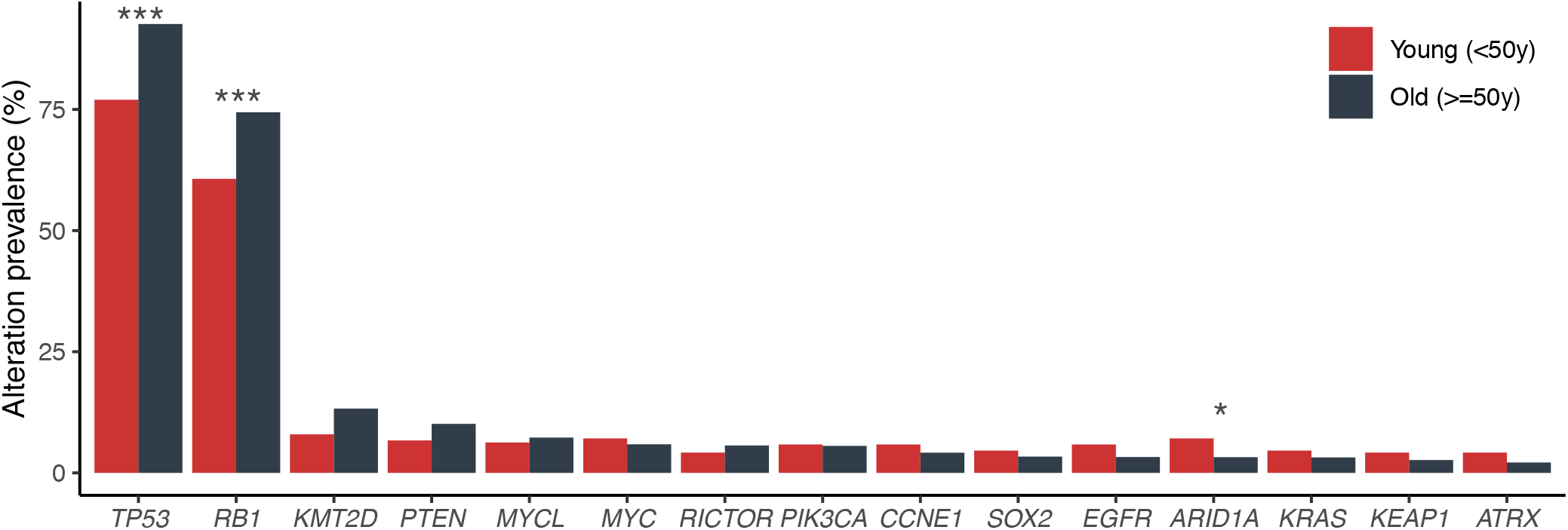
*TP53* and *RB1* mutations are more frequent in older patients with SCLC. Prevalence of genetic alterations in young (<50y) and older (50y+) patients with SCLC. Note the significantly higher frequency of *TP53* and *RB1* mutations in older patients, compared to young patients. Difference in alteration prevalence was assessed using a Fisher’s exact test with FDR-based correction (P value thresholds: *, 0.05; ***, 0.001).

**Figure S15.**
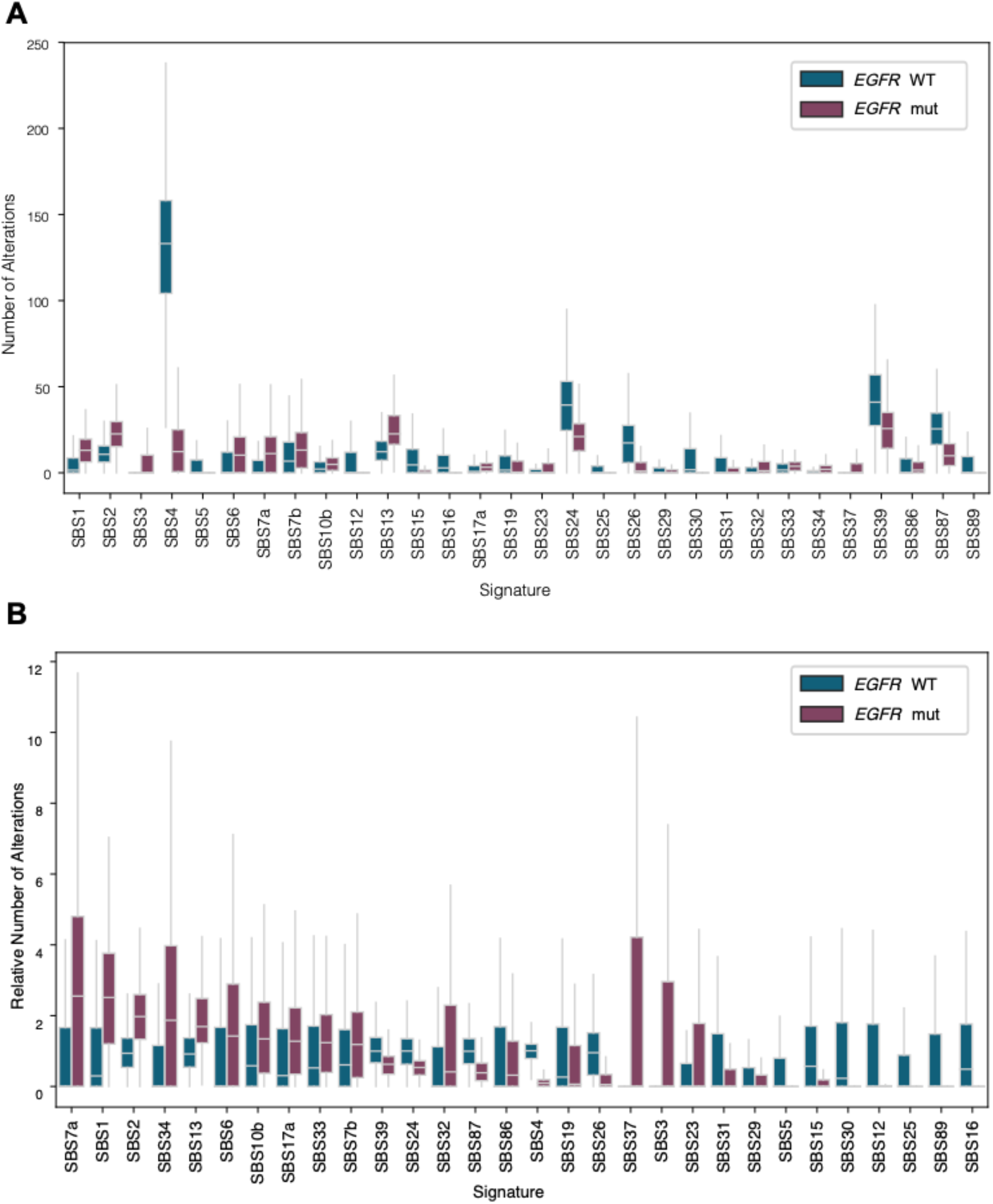
*EGFR*-mutant SCLC tumors exhibit unique mutational signature profiles. Analysis of the absolute number of alterations (**A**) and relative number of alterations (**B**) attributable to different single base substitution (SBS) signatures in *EGFR* wild-type and mutant tumors. The COSMIC annotations for the displayed mutational signatures include: SBS1 (clock-like), SBS2 (APOBEC), SBS3 (homologous recombination DNA damage repair), SBS4 (tobacco smoking), SBS5 (clock-like), SBS6 (DNA mismatch repair deficiency), SBS7a and SBS7b (ultraviolet light exposure), SBS10b (polymerase deficiency), SBS12 (unknown), SBS13 (APOBEC), SBS15 (DNA mismatch repair deficiency), SBS16, SBS17a, SBS19, SBS23 (unknown), SBS24 (aflatoxin exposure), SBS25 (chemotherapy treatment), SBS26 (DNA mismatch repair deficiency), SBS29 (tobacco chewing), SBS31 (platinum chemotherapy treatment), SBS32 (azathioprine treatment), SBS33, SBS34, SBS37, SBS39 (unknown), SBS86 (unknown chemotherapy treatment), SBS87 (thiopurine chemotherapy treatment), SBS89 (unknown).

**Figure S16:**
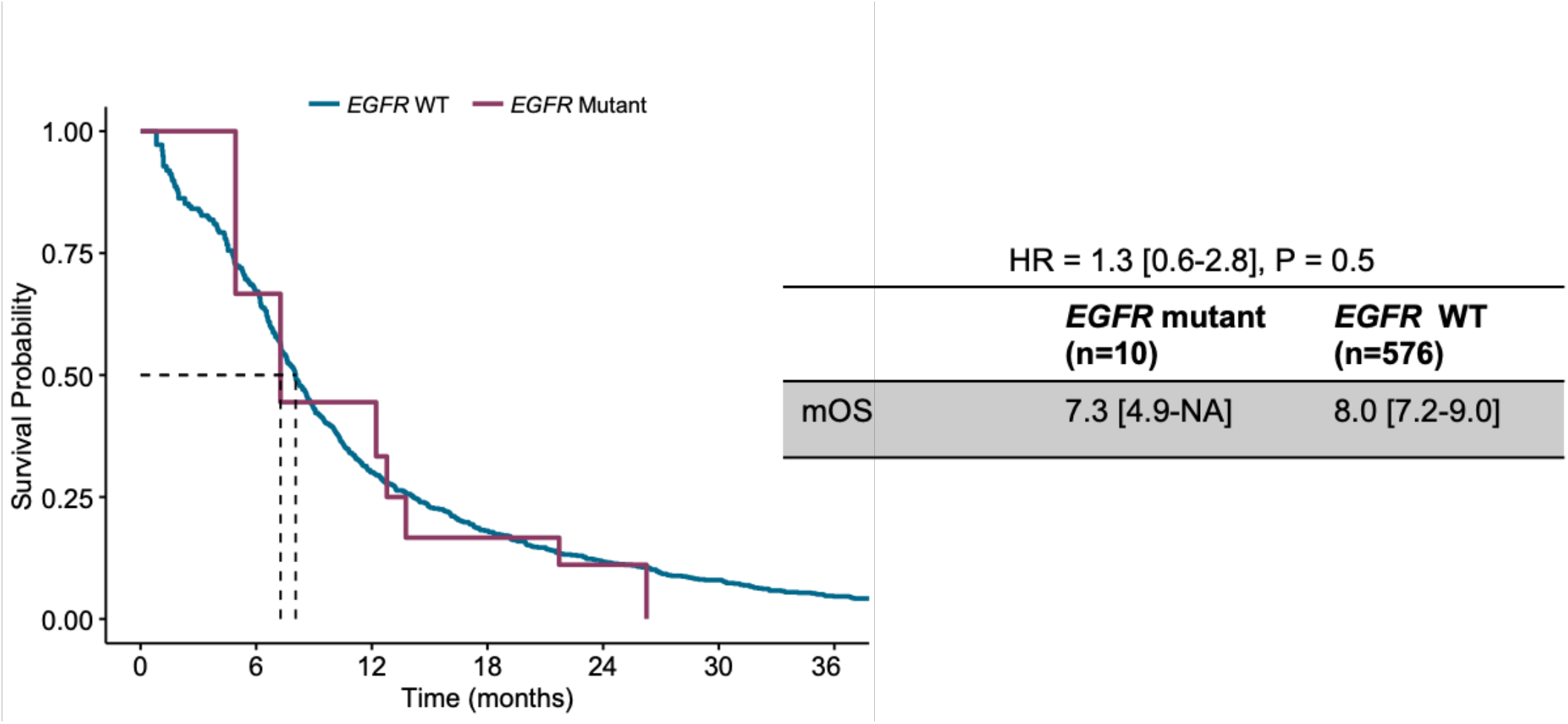
No overall survival differences based on *EGFR* mutation status in SCLC. Kaplan-Meir plots showing the overall survival in patients with SCLC tumors stratified by *EGFR* status (wild-type, WT or mutant).

**Figure S17:**
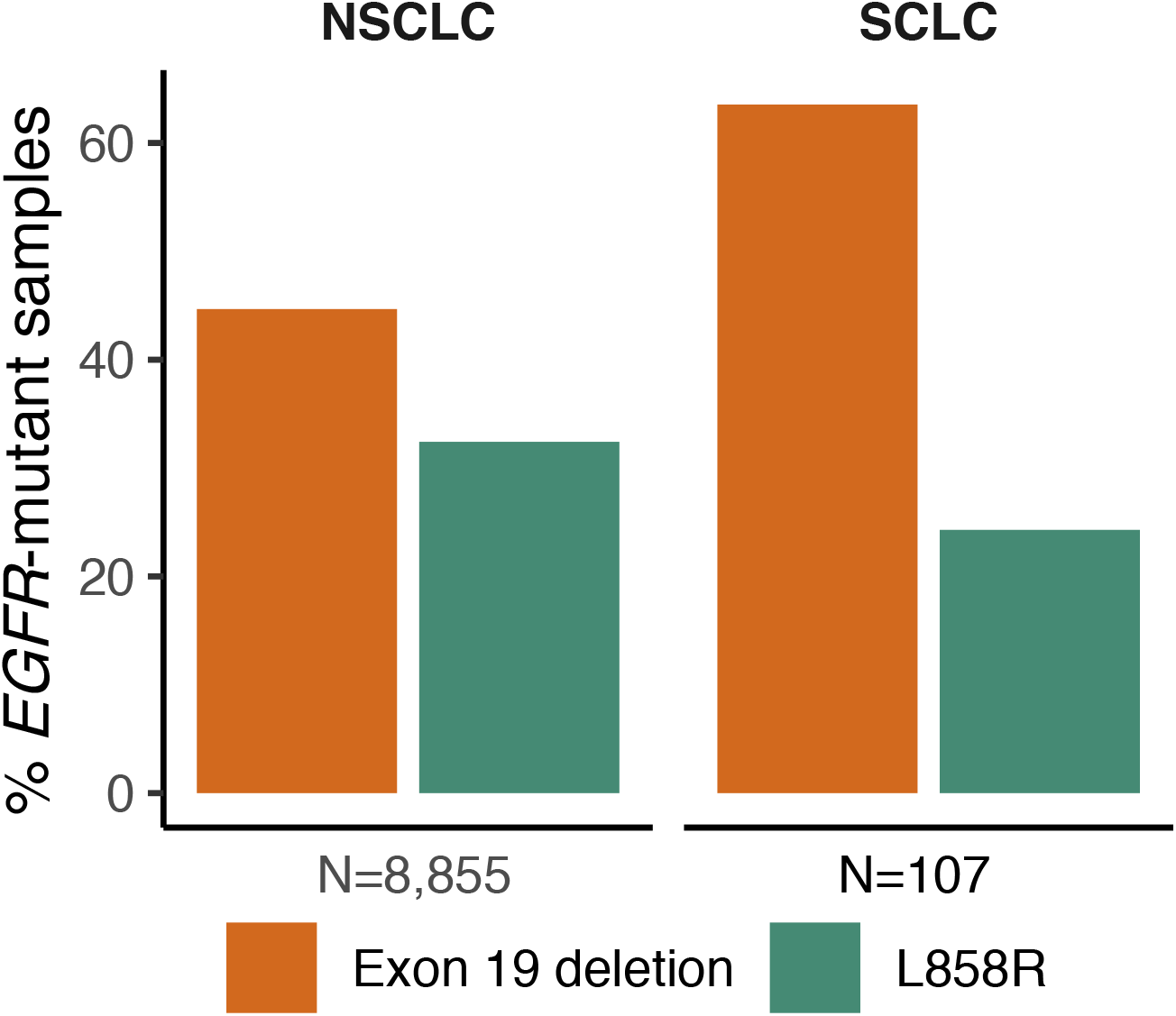
Putative transformed SCLC tumors from *EGFR*-mutant NSCLC are enriched for *EGFR* exon 19 deletions. Bar plots representing the proportion of *EGFR* L858R and *EGFR* exon 19-deleted tumors in the cohort of *EGFR*-altered SCLC in comparison to *EGFR*-altered NSCLC in the research dataset.

## SUPPLEMENTARY TABLES

See Excel file.

